# Nucleosome stability safeguards cell identity, stress resilience and healthy aging

**DOI:** 10.1101/2025.09.17.676776

**Authors:** Hiroshi Tanaka, Brenna S. McCauley, Clara Guida, Xue Lei, Sha Li, Tatiana M. Moreno, K’leigh Guillotte, Zong-Ming Chua, Adrianna Abele, Aashna Lamba, Rouven Arnold, Adarsh Rajesh, Marcos G. Teneche, Laurence Haddadin, Anagha Deshpande, Aniruddha J. Deshpande, Alexandre Colas, Caroline Kumsta, Michael Petrascheck, Rolf Bodmer, Weiwei Dang, Peter D. Adams

## Abstract

Nucleosomes are the minimal repeating units of chromatin. Their dynamic assembly and disassembly underpins chromatin organization and genome regulation. However, it remains unclear how intrinsic nucleosome stability contributes to higher-level yet fundamental cellular and organismal properties—such as preservation of cell identity, lineage specification, stress resilience and ultimately healthy aging. To address this, we tested the impact of decreased intrinsic nucleosome stability across multiple cell, tissue and organismal models by introducing histone mutants that weaken histone–histone interactions. While nucleosome instability did not broadly alter global chromatin accessibility, DNA damage, cell proliferation or viability, it impaired lineage-specific gene expression programs, altered lineage specification and activated intrinsic inflammatory and stress pathways in a manner reminiscent of aging in mouse tissues and human cells. Consistently, nucleosome instability accelerated the onset of age-associated transcriptional alterations and functional decline in *Caenorhabditis elegans* and *Drosophila melanogaster*, and reduced cellular resilience to exogenous perturbations— including environmental, epigenetic and mitotic stress—in human cells and *Saccharomyces cerevisiae*. These cross-species findings identify nucleosome stability as an evolutionarily conserved epigenetic safeguard that preserves cell identity and stress resilience and supports organismal function and healthy aging.

## Introduction

The capacity to maintain stable cell phenotypes under steady-state conditions—and particularly under stress—is fundamental to organismal longevity. Such phenotypic stability relies on cellular homeostatic mechanisms. Aging progressively erodes this homeostasis, compromising cell, tissue and organismal function and increasing susceptibility to degenerative conditions—including sarcopenia, fibrosis, cardiovascular disease, Alzheimer’s disease and cancer^1,2^. Chromatin serves as a central hub of cellular homeostasis by integrating intrinsic and extrinsic signals to orchestrate gene expression, DNA replication and repair, mitosis and genome maintenance, thereby governing cell fate and identity specification^3,4^. Thus, preserving chromatin integrity is essential for sustaining cellular phenotype and function over time. However, the homeostatic balance of chromatin—termed “chromostasis”^5,6^—is challenged during aging, resulting in global remodeling of chromatin structure, redistribution of histone and DNA modifications and changes in nucleosome positioning and density^7–10^. Emerging evidence suggests that such chromatin erosions are not merely consequences of aging, but actively drive the aging process^7,11^. This raises a fundamental question: to what extent does intrinsic chromatin stability preserve phenotypic fidelity and sustain organismal longevity throughout life?

Nucleosomes, the fundamental repeating units of chromatin, are semi-stable structures dynamically assembled and disassembled throughout a cell’s lifetime. Each nucleosome consists of ∼147 bp of DNA wrapped around a histone octamer composed of two copies each of H2A, H2B, H3 and H4, with structural stability dependent on specific histone– histone and histone–DNA interactions^12^. Although aging is associated with chromatin remodeling and nucleosome disorganization, it remains unclear the extent to which intrinsic nucleosome stability helps preserve cell identity and function, thereby supporting healthy aging. Notably, somatic mutations in cancer frequently affect specific histone residues—such as H2B D68 and E76—located at the H2B–H4 interface that stabilize the nucleosome core^13–17^. These observations suggest that decreased nucleosome stability *per se* may compromise the fidelity of gene regulation, erode cell identity, increase susceptibility to stress and ultimately threaten organismal healthspan. To test this hypothesis, we introduced defined histone mutants that decrease intrinsic nucleosome stability across multiple cell, tissue and organismal models. This enabled direct interrogation of the role of nucleosome stability in chromatin regulation, phenotypic robustness and healthy aging.

## Results

### Characterization of nucleosome-destabilizing histone mutants

As tools to probe the impact of nucleosome instability, we employed histone H2B mutants that disrupt histone–histone interactions within the nucleosome. The D68 residue of H2B forms hydrogen bonds with K91, T96 and Y98 of one H4 molecule, while E76 interacts with Y72 and R92 of the other H4 (**Fig. 1a**). Hence, mutations at these residues within H2B disrupt H2B– H4 interactions and confer nucleosome instability^13,15,18^. We transduced immortalized IMR-90 human fetal lung fibroblasts (IMR-90 SV40-T) with lentiviral vectors expressing either wild-type (WT) or mutant H2B (D68N, E76K, E76R or double mutant D68N/E76K) (**Extended Data Fig. 1a–c**). Although the E76K and E76R substitutions both replace acidic E with basic K or R, the latter is more basic and bulkier and thus predicted to be more disruptive of nucleosome structure. Expression levels of H2B D68N and E76K were comparable to those of WT and similar to endogenous H2B, whereas E76R and D68N/E76K showed slightly lower expression, possibly linked to their predicted heightened nucleosome instability. Expression of these mutants did not cause overt defects in cell proliferation or steady-state abundance of γH2AX, a marker of DNA damage (**Extended Data Fig. 1d,e**). EGFP-tagged WT and mutant H2B localized to the nucleus and were fully incorporated into mitotic chromosomes (**Extended Data Fig. 1f,g**), confirming their incorporation into chromatin. However, although sequential salt fractionation revealed that both WT and mutant H2B predominantly localized in the chromatin fraction, the mutants were extracted at a lower salt concentration (0.9 M NaCl) compared to endogenous H2B and ectopic H2B WT (1.2 M NaCl), indicating weaker chromatin association (**Extended Data Fig. 1h**). Consistently, fluorescence recovery after photobleaching (FRAP) demonstrated increased mobility of mutant proteins in the nucleus compared to WT (**Fig. 1b**). As predicted, E76R and D68N/E76K conferred higher mobility compared to E76K. Using H2A-EGFP, we also observed increased H2A mobility in the presence of H2B mutants, consistent with the predicted destabilization of the H2A–H2B dimer in chromatin (**Extended Data Fig. 1i**). Complementing these findings, pulse-chase labeling of SNAP–tagged H2B showed faster turnover of mutant proteins compared to WT (**Fig. 1c and Extended Data Fig. 1j,k**). These results indicate that although both WT and mutant H2B are similarly incorporated into chromatin, the mutants induce nucleosome instability in cells.

**Fig. 1.**
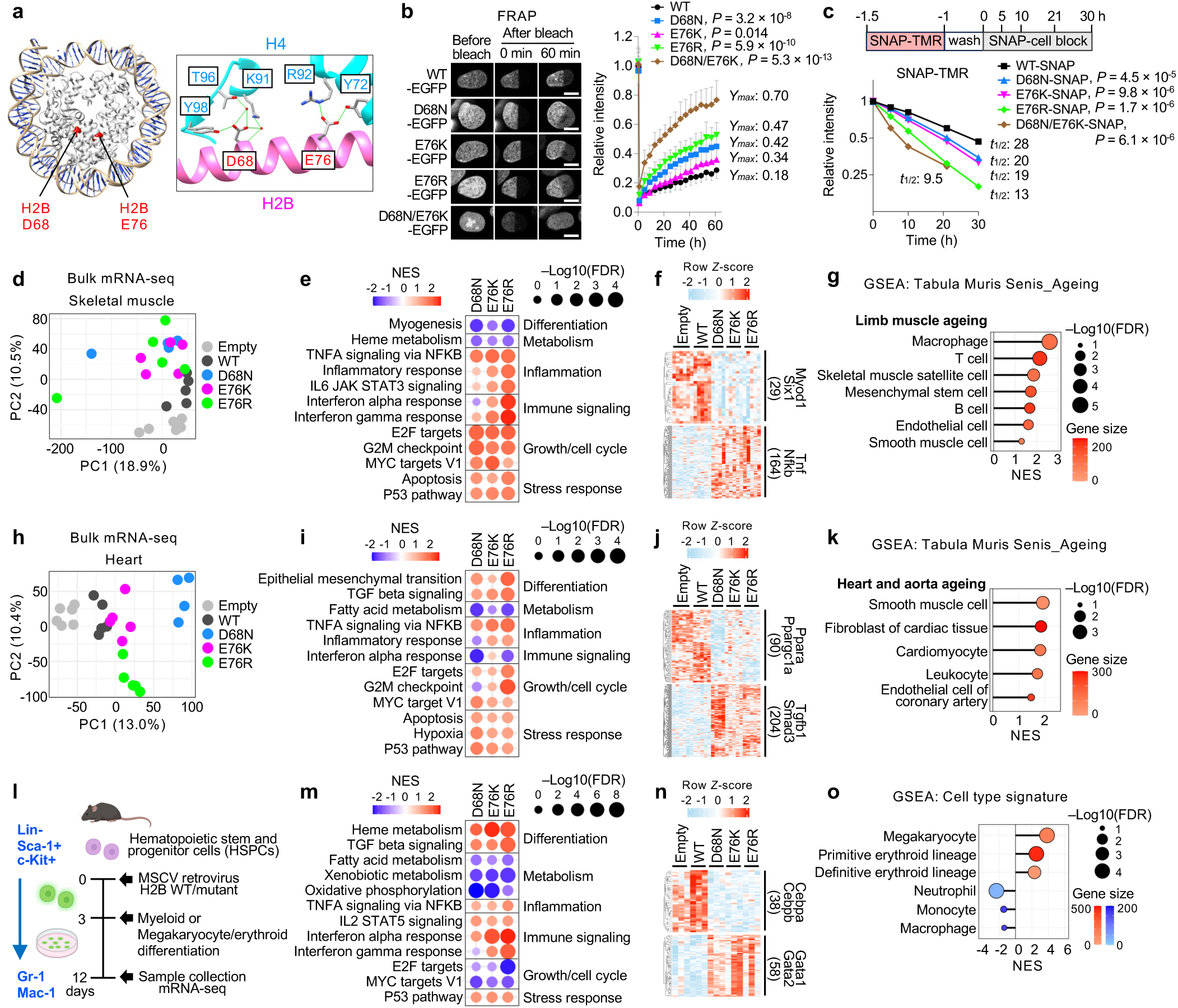
Nucleosome instability compromises lineage-specific gene expression programs and induces inflammatory, immune and stress responses, recapitulating aging-like signatures across mouse tissues. **a,** Structural location of histone H2B residues D68 and E76 mutated in this study. H2B is shown in magenta, H4 in cyan. **b,** Nucleosome stability assessed by FRAP in IMR-90 SV40-T cells expressing EGFP-tagged H2B WT or mutants. Scale bar, 10 μm. Relative intensity was calculated by comparing the bleached site to the unbleached site. Data represent mean ± s.d. from >15 nuclei. The plateau value (*Ymax*) was calculated from the fitted recovery curve. *P*-values were calculated using unpaired, two-tailed Student’s *t*-tests comparing each mutant to WT at the final time point. **c,** Histone turnover in live cells assessed using SNAP-tagged H2B WT or mutants labeled with SNAP-TMR. Data represent mean ± s.d. of total field fluorescence from >2600 cells across three independent wells per condition. Half-life (*t*_1/2_) was calculated from the fitted decay curve. *P*-values were calculated using unpaired, two-tailed Student’s *t*-tests comparing each mutant to WT at the final time point. **d,** PCA of bulk mRNA-seq from skeletal muscle expressing H2B WT, mutants or empty vector. Each dot represents an individual mouse (n = 6, Empty; 5, WT; 4, D68N; 5, E76K; 5, E76R). **e,** GSEA of hallmark pathways in transduced skeletal muscle comparing each mutant to Empty and WT. (See Extended Data Fig. 3c for full pathways). **f,** IPA upstream regulator analysis showing suppression of Myod1- or Six1-associated genes and activation of Tnf- or Nfkb-associated genes across mutant-expressing skeletal muscle compared to Empty and WT. **g,** GSEA of Tabula Muris Senis aging signatures in transduced skeletal muscle comparing all mutants to Empty and WT. **h,** PCA of bulk mRNA-seq from heart expressing H2B WT, mutants or an empty vector. Each dot represents an individual mouse (n = 6, Empty; 5, WT; 4, D68N; 5, E76K; 5, E76R). **i,** GSEA of hallmark pathways in transduced heart comparing each mutant to Empty and WT. (See Extended Data Fig. 3j for full pathways). **j,** IPA upstream regulator analysis showing suppression of Ppara- or Ppargc1a-associated genes and activation of Tgfb1- or Smad3-associated genes across mutant-expressing heart compared to Empty and WT. **k,** GSEA of Tabula Muris Senis aging signatures in transduced heart comparing all mutants to Empty and WT. **l,** Schematic of experimental design. MSCV-based retroviral vectors expressing H2B WT or mutants were introduced into HSPCs (Lin⁻Sca-1⁺c-Kit⁺), and myeloid or megakaryocyte/erythroid differentiation was assessed. **m,** GSEA of hallmark pathways in transduced HSPCs comparing each mutant to Empty and WT. (See Extended Data Fig. 4e for full pathways). **n,** IPA upstream regulator analysis showing suppression of Cebpa- or Cebpb-associated genes (myeloid regulators) and activation of Gata1- or Gata2-associated genes (megakaryocyte/erythroid regulators) in mutant-expressing HSPCs 9 days after differentiation. **o,** GSEA of cell type signatures indicating increased megakaryocyte and erythroid lineage features and decreased myeloid (neutrophil, monocyte and macrophage) features in mutant-expressing cells compared to Empty and WT.

### Nucleosome instability impairs lineage-specific gene expression programs and induces inflammatory, stress and aging-like signatures

To examine the *in vivo* impact of nucleosome instability, we retro-orbitally injected AAV9 vectors encoding H2B WT or mutants (D68N, E76K or E76R) under the control of the CMV promoter into 3-week-old C57BL/6J mice (**Extended Data Fig. 2a**). Four weeks later, body weight and gross appearance were unaffected, and expression of WT and mutant H2B was detected in skeletal muscle, heart and liver, with the expression levels <50% of endogenous H2B (**Extended Data Fig. 2b–e**, see total H2B Western blot in 2c). Transcriptomic analysis revealed modest transcriptional changes in liver, but skeletal muscle and heart each exhibited over 1,300 genes either upregulated or downregulated, mostly tissue specific (**Extended Data Fig. 3a,b**). In skeletal muscle, principal component analysis (PCA) separated mutants from controls (Empty and WT) (**Fig. 1d**), and gene set enrichment analysis (GSEA) showed repression of myogenesis and heme metabolism genes, alongside activation of inflammatory, immune, growth and stress-response genes in mutant-expressing muscles (**Fig. 1e and Extended Data Fig. 3c**). Ingenuity Pathway Analysis (IPA) inferred suppression of Myod1- and Six1-regulated myogenic programs, including genes involved in muscle contraction and glycolysis, while activating inflammatory, immune and stress pathways driven by Tnf, Nfkb, Cebpb, Il6, Stat3, Ifng, Stat1, Hif1a, Nfe2l2, Myc and p53 (**Fig. 1f and Extended Data Fig. 3d,e**). In line with immune activation, chemokines *Ccl22*, *Ccl12* and *Cxcl10*, known to recruit macrophages and T cells and accumulate during aging and regeneration^19,20^, were upregulated, accompanied by enrichment of proinflammatory M1 macrophage transcriptional traits (**Extended Data Fig. 3f,g**). Upregulation of p53 target genes (*Bbc3/Puma*, *Bax* and *Cdkn1a/p21*), muscle regeneration-associated genes (*Tceal7*, *H19* and *Pim1*)^21–23^ and Myc target ribosomal protein genes (RPLs and RPSs) suggests maladaptive regenerative responses reminiscent of aged muscle^24^ (**Extended Data Fig. 3h,i**). Cell type signature profiles using the mouse aging cell atlas, *Tabula Muris Senis*^25^, revealed that mutant-expressing skeletal muscle recapitulated aging-like transcriptional signatures across macrophages, T cells, muscle satellite cells, mesenchymal stem cells and B cells (**Fig. 1g**).

In heart, mutants repressed fatty acid metabolism genes, while activating epithelial mesenchymal transition (EMT), TGFβ signaling, inflammation, growth and stress-response genes (**Fig. 1h,i and Extended Data Fig. 3j**). IPA predicted suppression of Ppara and Ppargc1a, master regulators of fatty acid metabolism^26^, while activating the TGFβ1–Smad3 axis, a central mediator of EMT and myogenic processes^27^, as well as inflammatory, immune and stress pathways driven by Tnf, Cebpb, Il4, Stat3, Hif1a, Nfe2l2, Vegfa, Myc and p53 (**Fig. 1j and Extended Data Fig. 3k,l**). Unlike skeletal muscle, mutant hearts upregulated the anti-inflammatory cytokine *Ccl17*, which plays a critical role in heart failure and cardiovascular aging^28^ (**Extended Data Fig. 3m**), suggesting tissue-specific immune adaptation to nucleosome instability. Activation of proliferative and growth programs, including ribosomal protein genes, suggested regenerative activity or hypertrophic growth responses in the heart (**Extended Data Fig. 3n**). Cell type signature profiles revealed patterns reminiscent of aging across smooth muscle cells, fibroblasts, cardiomyocytes, leucocytes and endothelial cells (**Fig. 1k**). Collectively, these results demonstrate that nucleosome instability compromises lineage-specific gene expression programs, triggers persistent inflammatory, immune and p53-associated stress responses, and recapitulates aging-like signatures across multiple tissues.

### Nucleosome instability compromises cell-intrinsic lineage specification

Given that nucleosome instability perturbs lineage-specific gene expression programs and recapitulates aging-like features in mouse tissues, we next examined its direct impact on cell-intrinsic differentiation potential using mouse hematopoietic stem and progenitor cells (HSPCs). Hematopoiesis is a dynamic process whereby HSPCs commit to specific blood cell lineages in response to differentiation cues^29,30^, and aging reduces this capacity while skewing output toward myeloid and megakaryocyte/erythroid lineages^31–34^. To test whether nucleosome instability affects HSPC differentiation, freshly isolated mouse HSPCs (Lin^-^Sca-1^+^c-Kit^+^) were transduced with MSCV retroviral vectors expressing H2B WT or mutants and cultured in semi-solid methylcellulose media containing cytokines and hormones that promote myeloid and megakaryocyte/erythroid differentiation (**Fig. 1l and Extended Data Fig. 4a**). After 8 days, three colony types were detected: colony-forming unit granulocyte (CFU-G), macrophage (CFU-M) and granulocyte/macrophage bipotent progenitors (CFU-GM) (**Extended Data Fig. 4b**). Notably, H2B mutants markedly reduced CFU-GM formation (**Extended Data Fig. 4c**). Flow cytometry confirmed retention of Sca-1^+^c-Kit^+^ cells, with reduced Gr-1^+^ and Mac-1^+^ populations (**Extended Data Fig. 4d**), indicating impaired differentiation of granulocytic and macrophage lineages, with persistence of undifferentiated progenitors.

Transcriptomic profiling revealed downregulation of fatty acid metabolism, xenobiotic metabolism and oxidative phosphorylation genes, while upregulating heme metabolism, TGFβ signaling, inflammation, interferon and p53 pathway genes (**Fig. 1m and Extended Data Fig. 4e**). IPA indicated suppression of Cebpa and Cebpb—key drivers of myeloid specification^35,36^—while activating Gata1, Gata2, Runx1, Nfe2, Fli1 and Stat5b, which promote heme metabolism and megakaryocyte/erythroid differentiation^37–39^ (**Fig. 1n and Extended Data Fig. 4f**). Consistently, cell type signature profiles indicated loss of neutrophil, monocyte and macrophage signatures, with enrichment of megakaryocyte and erythroid signatures (**Fig. 1o**). This skewed output was accompanied by activation of Tnf, Nfkb, Irf3/7/9, Ifng, Stat1 and p53 pathways—known to impair HSPC differentiation and promote lineage bias^40–42^ (**Extended Data Fig. 4g**). Together, these data demonstrate that nucleosome instability compromises cell-intrinsic differentiation potential of HSPCs, impairing lineage-determining transcriptional programs and skewing transcriptional output toward megakaryocyte/erythroid lineages, thereby partially recapitulating signatures of hematopoietic aging.

### Nucleosome instability drives aging-like stress–identity convergence

To further assess the cell-intrinsic impact of nucleosome instability, we profiled the transcriptome of IMR-90 SV40-T human fibroblasts expressing H2B WT or mutants, alongside unmodified primary human dermal fibroblasts from young (ages 23–33) and aged (ages 64– 72) donors. PCA clearly separated mutant-expressing cells from controls (Empty and WT), and aged from young fibroblasts (**Fig. 2a,b**). GSEA revealed that H2B mutants recapitulated aged fibroblast signatures, including upregulation of myogenesis, EMT, TGFβ signaling, inflammatory and stress-response programs (**Fig. 2c and Extended Data Fig. 5a**). Despite these phenotypic similarities, mutant-expressing cells exhibited distinct cell-cycle and growth profiles compared to aged fibroblasts, indicating that similarities between H2B mutants and aging are uncoupled from proliferative state. IPA predicted activation of SRF, MRTFA/B and SMAD3—key regulators of smooth muscle identity and myofibroblast differentiation^43–45—^ together with proinflammatory and stress pathways, including TNF, NFKB, STAT3, AP1, MYC, FLT1, HIF1A, NFE2L2 and p53 pathways (**Fig. 2d,e and Extended Data Fig. 5b**). The magnitude of these responses increased as nucleosome stability decreased, consistent with greater effects at higher destabilization (**Fig. 2f**). Together with mouse data, these results demonstrate that nucleosome instability causally drives an aging-like phenotypic shift in human fibroblasts.

**Fig. 2.**
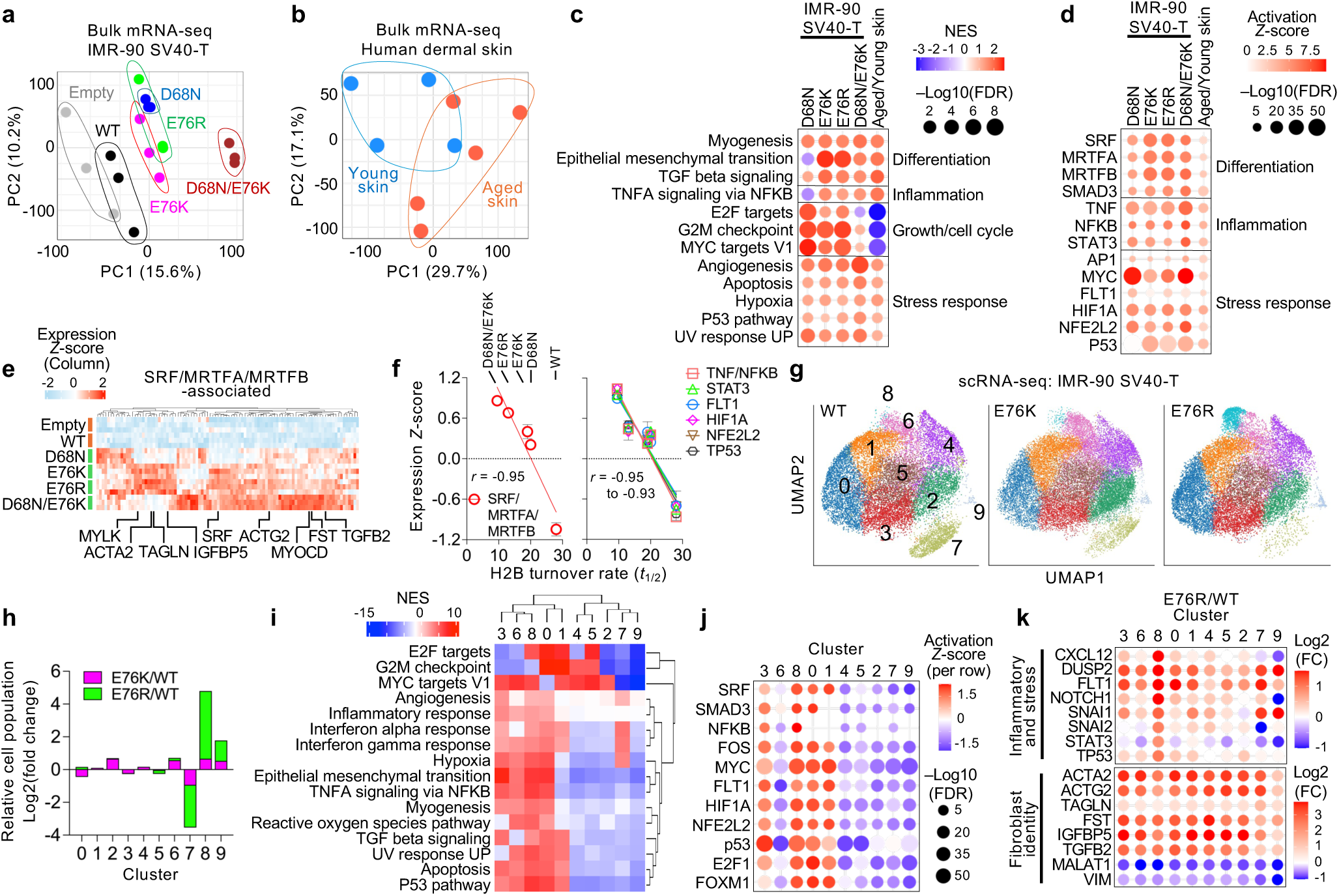
Nucleosome instability induces aging-like stress–identity convergence in single cells. **a**, PCA of bulk mRNA-seq from IMR-90 SV40-T cells expressing H2B WT, mutants or empty vector. Each dot represents an individual replicate (n = 3 per condition). **b**, PCA of bulk mRNA-seq from dermal skin biopsies comparing young (ages 23–33, n = 4) and aged (ages 64–72, n = 5) individuals. **c**, GSEA of hallmark pathways in mutant-expressing IMR-90 SV40-T cells compared to Empty and WT cells and in aged primary skin fibroblasts compared to young fibroblasts. See Extended Data Fig. 5a for full pathways. **d**, IPA upstream regulator analysis showing activation of differentiation, inflammation and stress-response pathways in mutant-expressing IMR-90 SV40-T cells compared to Empty and WT cells and in aged primary skin fibroblasts compared to young fibroblasts. **e**, Expression *Z*-scores of 98 SRF/MRTFA/MRTFB-associated genes upregulated in mutant-expressing IMR-90 SV40-T cells compared to Empty and WT cells. **f**, Correlation between H2B turnover rates (*t*_1/2_, measured in Fig. 1c) and mean expression *Z*-scores of gene sets associated with SRF/MRTFA/MRTFB, TNF/NFKB, STAT3, FLT1, HIF1A, NFE2L2 and TP53 that are upregulated in mutant-expressing IMR-90 SV40-T cells compared to Empty and WT cells. Each point represents a mutant or WT condition. Linear regression lines and Pearson correlation coefficients (*r*) are shown. **g**, UMAP plot of scRNA-seq from IMR-90 SV40-T cells expressing H2B WT, E76K or E76R. **h**, Log_2_ fold-changes in cell population abundance across clusters for E76K/WT and E76R/WT comparisons in scRNA-seq analysis. **i**, GSEA of hallmark pathways enriched in each cluster compared to all other clusters in scRNA-seq analysis. See Extended Data Fig. 5e for full results. **j**, IPA upstream regulator analysis showing representative signaling pathways enriched in each cluster compared to all other clusters. **k**, Cluster-wise log_2_ fold-changes of inflammatory/stress-related or fibroblast identity gene expression in scRNA-seq analysis of E76R-versus WT-expressing IMR-90 SV40-T cells.

Building on these bulk mRNA-seq observations, we next applied single-cell RNA-seq (scRNA-seq) to IMR-90 SV40-T cells expressing H2B WT, E76K or E76R in order to dissect stress and identity programs at single-cell resolution. UMAP analysis identified nine transcriptionally distinct clusters (**Fig. 2g**). Cells expressing the mutants notably expanded clusters 8 and 9 while reducing cluster 7 (**Fig. 2h and Extended Data Fig. 5c**), indicative of altered cell-state distribution. Pathway analysis revealed that cluster 8 was enriched for both myogenesis and inflammatory/stress-response programs, whereas clusters 7 and 9 exhibited fewer enriched pathways (**Fig. 2i and Extended Data Fig. 5d,e**). IPA confirmed concurrent activation of stress and SRF-regulatory pathways in cluster 8 (**Fig. 2j and Extended Data Fig. 5f**). Differential expression analysis between WT and mutants demonstrated consistent upregulation of inflammatory/stress-response genes across clusters, most prominently in cluster 8 (**Fig. 2k and Extended Data Fig. 5g**). Meanwhile, SRF/MRTFA/B-associated genes, such as ACTA2 and ACTG2, were broadly induced across clusters, accompanied by suppression of mesenchymal markers *MALAT1* and *VIM*, indicating a global basal-level shift toward a myofibroblast-like identity (Fig. 2k). Together, these results demonstrate that nucleosome instability reshapes the cellular landscape by promoting convergence of stress and identity/myogenic programs, altering cluster composition and inducing a myofibroblast-like shift reminiscent of aged fibroblasts^46,47^.

### Nucleosome instability alters transcriptional programs beyond chromatin accessibility

To assess how nucleosome instability affects gene expression, we profiled IMR-90 SV40-T cells expressing H2B WT or mutants using ATAC-seq and SLAM-seq. MNase digestion showed no measurable change in global chromatin accessibility (**Extended Data Fig. 6a**). To examine locus-specific effects, we performed ATAC-seq with TGFβ1 treatment as a positive control (**Extended Data Fig. 6b**). E76K expression induced 204 open and 69 closed regions, with more than half of the open regions overlapping TGFβ-responsive sites (126/204) (**Extended Data Fig. 6c**), consistent with increased expression of TGFβ pathway genes (**Fig. 2c**). By contrast, TGFβ treatment induced more than 8,000 open or closed regions in both E76K and control cells. Moreover, E76K-responsive sites were largely confined to distal regulatory elements (>±5 kb, up to 100 kb from TSS) (**Extended Data Fig. 6d**), indicating that nucleosome instability has only minimal impact on promoter- or gene-associated steady-state accessibility.

To further assess the link between H2B mutant-induced chromatin accessibility and transcriptional changes, we performed SLAM-seq to detect nascent transcripts (**Extended Data Fig. 6e**). SLAM-seq revealed transcriptional activation of myogenic and stress-related pathway genes, including SRF-associated myofibroblast-related genes such as *ACTA2* and *ACTG2*, similar to those observed by bulk mRNA-seq (**Extended Data Fig. 6f**). Strikingly, most transcriptional changes detected by SLAM-seq occurred independently of detectable changes in chromatin accessibility (**Extended Data Fig. 6g.h**). Thus, nucleosome instability drives transcriptional reprograming linked to increased nucleosome destabilization and turnover, but not to detectable changes in global or local steady-state chromatin accessibility.

### Nucleosome instability drives premature aging

Since nucleosome instability elicits aging-like features in mouse tissues and human cells, we examined its organismal impact in *Caenorhabditis elegans* and *Drosophila melanogaster*. Histone H2B is highly conserved among worms, flies and human, with human E76 corresponding to E73 in worms and flies (**Extended Data Fig. 7a**). Using CRISPR/Cas9, we engineered *C. elegans* strains to express E73K or E73R mutant H2B from *his-41/C50F4.5* locus^48^ (**Fig. 3a and Extended Data Fig. 7b**). Notably, the E73R mutant significantly shortened lifespan by 19% compared to N2 control, whereas the less disruptive E73K mutant had no discernible effect (**Fig. 3b**). Transcriptomic profiling at post-adult day 3 revealed that E73R mutants showed distinct transcriptomic signatures from WT, including suppression of embryonic development and cell cycle genes, and activation of stress-response pathways (**Fig. 3c and Extended Data Fig. 7c–c**). By day 10, defense response genes, including immune components, were markedly diminished (**Extended Data Fig. 7h,i**). Of the 2,396 genes downregulated in E73R mutants at day 3, 1,089 also declined during normal aging in WT worms by day 10 (**Fig. 3d,e**). These genes were enriched for embryo development and cell cycle regulation (**Fig. 3f**), suggesting accelerated premature decline of the germline which comprises a large proportion of total cells in adult worms. Conversely, E73R upregulated 2,646 genes at day 3, including 645 genes upregulated during WT aging (**Fig. 3d,g**). These were enriched for pathways involving cellular senescence, organelle localization, locomotor activity and stress responses, reflecting early onset of degenerative phenotypes (**Fig. 3h**).

**Fig. 3.**
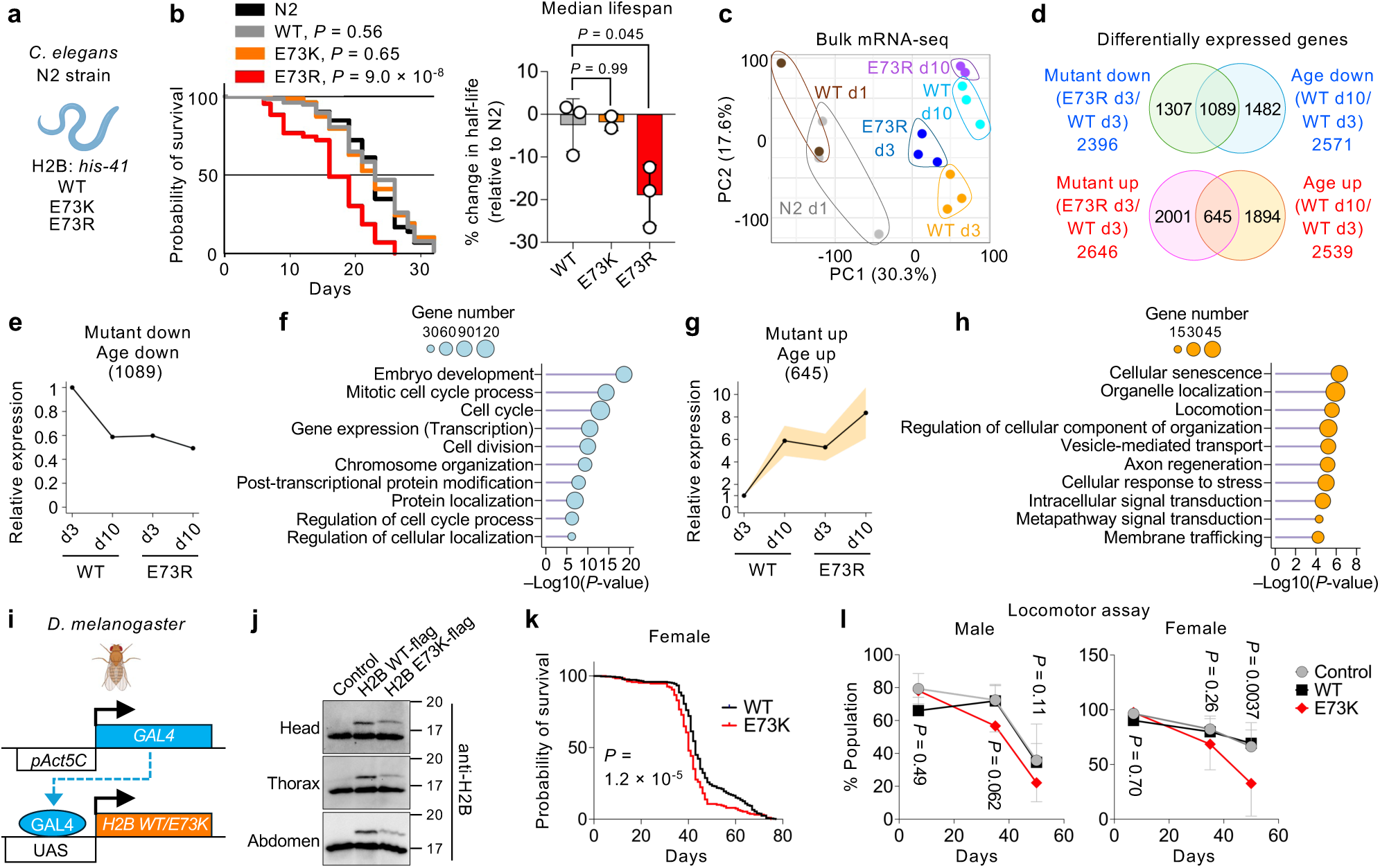
Nucleosome instability accelerates aging phenotypes in C. elegans and D. melanogaster. **a**, Schematic model of *C. elegans* expressing H2B/*his-41* WT or mutant generated via CRISPR-Cas9 in the N2 strain background. **b**, Representative Kaplan–Meier survival curves of *C. elegans* expressing H2B WT, E73K, E73R or unmodified N2 strain from three independent experiments. Log-rank tests were used to compare WT or each mutant to N2; *P*-values are indicated on the graph. Bar plot depicts median lifespan relative to N2 across three independent experiments. Statistical significance among groups was assessed by one-way ANOVA (*P* = 0.034), followed by post hoc pairwise comparisons using estimated marginal means (emmeans) with Tukey’s HSD adjustment. Only Tukey-adjusted *P*-values are shown. **c**, PCA of transcriptomic profiles from adult *C. elegans* at days 1, 3 and 10 post-adulthood. Each dot represents an individual worm (n = 3, N2 d1; 2, WT d1; 3, WT d3; 3, E73R d3; 3, WT d10; 2, E73R d10). **d**, Differentially expressed genes from mRNA-seq in *C. elegans* expressing H2B WT or E73R at days 3 or 10. **e**, Relative expression of genes downregulated both by E73R at day3 (E73R d3 vs WT d3, mutant down) and by aging in WT (WT d10 vs WT d3, age down). Shaded areas indicate 95% confidence intervals (sky blue, though nearly invisible due to low values). **f**, Gene ontology enrichment (Metascape) of 1,089 genes downregulated by both E73R and with age. **g**, Relative expression of genes upregulated both by E73R at day3 (E73R d3 vs WT d3, mutant up) and by aging in WT (WT d10 vs WT d3, age up). Shaded areas indicate 95% confidence intervals (orange). **h**, Gene ontology enrichment (Metascape) of 645 genes upregulated with both E73R and with age. **i**, Schematic model of the GAL4-UAS system used in *D. melanogaster* to express H2B WT or E73K under the control of the *Act5C* ubiquitous promoter. **j**, Western blot of H2B-Flag in dissected fly tissues. **k**, Representative Kaplan–Meier survival curves of female *D. melanogaster* expressing H2B WT or E73K from two independent experiments. A total of 365 WT and 188 E73K flies were analyzed. *P*-value was calculated using the log-rank test comparing E73K to WT. **l**, Locomotor performance assay of male and female *D. melanogaster* expressing H2B WT, E73K or wild-type controls. Data were analyzed by two-way ANOVA. In females, both genotype and age effects were significant (*P* = 0.030 and 1.1 × 10^−4^, respectively), whereas in males only age was significant (*P* = 7.1 × 10^−8^). Post-hoc pairwise comparisons between E73K and Control/WT at each age were calculated using emmeans with Tukey’s HSD adjustment; comparisons are shown for both sexes for consistency, and only Tukey-adjusted *P*-values are shown.

In *D. melanogaster*, we expressed H2B mutant using the GAL4/UAS system under the control of the ubiquitously active *Act5C* promoter (**Fig. 3i**). Western blotting confirmed expression of ectopic H2B WT and E73K across whole bodies, at levels below that of endogenous H2B (**Fig. 3j**). Notably, E73K expression modestly reduced lifespan in female flies, but had no measurable effect in males (**Fig. 3k and Extended Data Fig. 7j**). Consistently, the mutants accelerated the age-associated decline in locomotor performance in females, with males also showing a subtle trend toward reduction (**Fig. 3l**). Collectively, these findings demonstrate that nucleosome instability promotes premature aging phenotypes, from transcriptome to function to lifespan, across evolutionarily diverse organisms.

### Nucleosome instability reduces stress resilience

Nucleosome-destabilizing mutants consistently upregulated stress-response genes across models, suggesting decreased cellular stress resilience. Indeed, IMR-90 SV40-T cells expressing these mutants exhibited reduced viability following heat stress or irradiation (**Fig. 4a**). To assess evolutionary conservation, we tested stress tolerance in *Saccharomyces cerevisiae*. As yeast possess two H2B genes (*HTB1* and *HTB2*), we replaced *HTB1* with mutants equivalent to human D68N, E76K and E76R (*HTB1* D71N, E79K or E79R) and deleted *HTB2* (**Extended Data Fig. 8a,b**). As in human fibroblasts, these mutants did not impair proliferation (**Extended Data Fig. 8c**) or replicative lifespan (**Extended Data Fig. 8d**), but markedly reduced tolerance to cold, heat, UV and genotoxic agents such as camptothecin (CPT) and methyl methanesulfonate (MMS) (**Fig. 4b**).

**Fig. 4.**
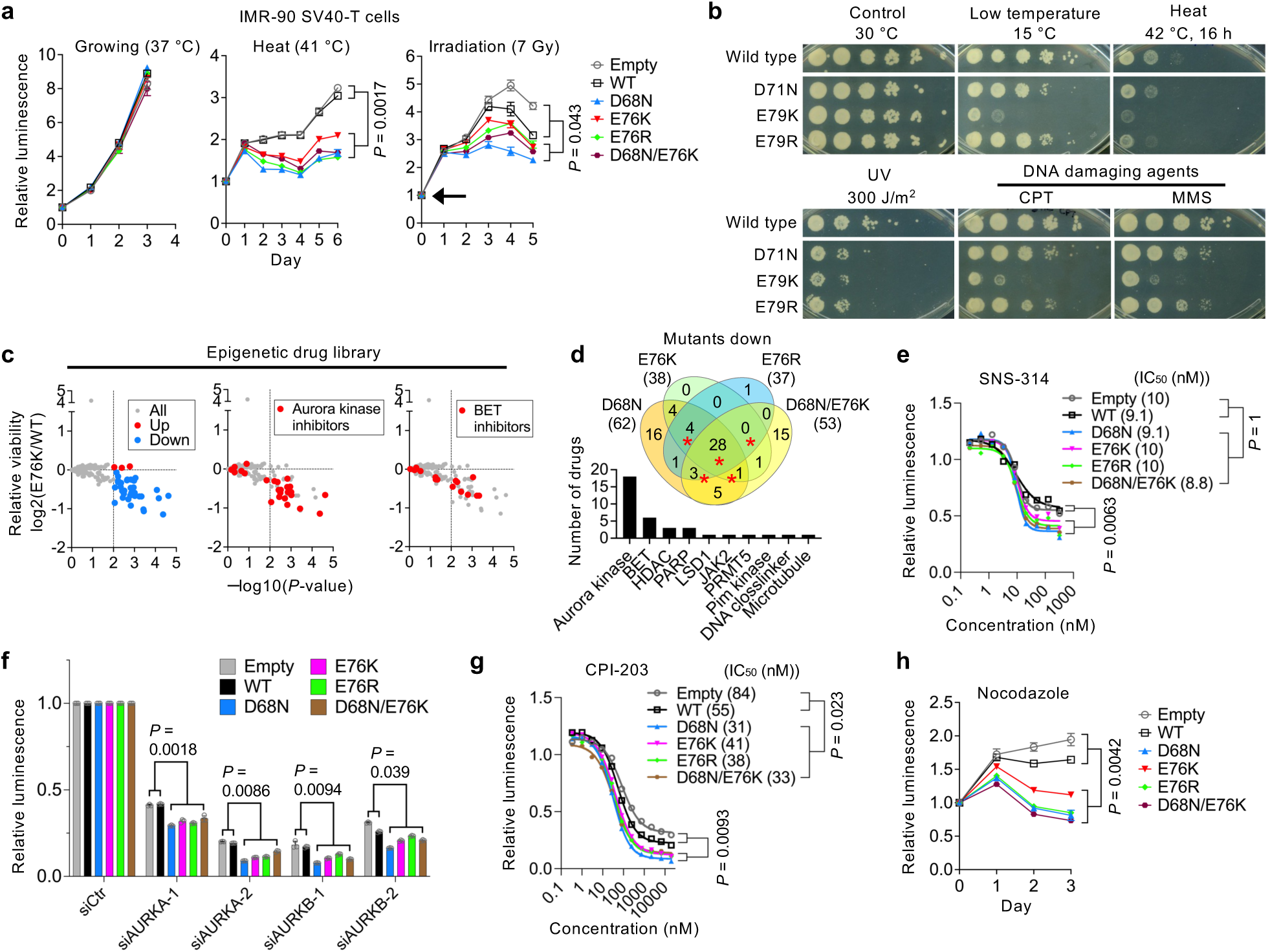
Nucleosome instability reduces epigenetic resilience and heightens vulnerability to exogenous perturbations. **a**, Cell viability assay under normal condition (37 °C), heat stress (41 °C) or after irradiation (7 Gy) in IMR-90 SV40-T cells expressing H2B WT, mutants or an empty vector. Arrow indicates the day of irradiation. Data represent mean ± s.d. from three replicates, each consisting of 4–10 wells per condition. *P*-values were calculated using unpaired, two-tailed Student’s *t*-tests comparing the means of all mutants to Empty and WT at the final time point. Data shown are representative of two independent experiments for growing and heat stress, and similar trends were observed with two independent experiments at 5 Gy and 10 Gy irradiation (data not shown). **b**, Stress resistance assays in yeast strains expressing HTB1 WT, D71N, E79K or E79R under various stress conditions: control (30 °C for 2 days), low temperature (15 °C for 5 days), heat shock (42 °C for 16 h, then 30 °C until 2 days), UV irradiation (300 J/m², then 30 °C until 2 days) and DNA damage (camptothecin (CPT) or methyl methanesulfonate (MMS) for 2 days). Cells were spotted as 1:10 serial dilutions starting from 2 OD_600_ from left to right. Experiments were performed for 3, 4 and 3 independent clones for D71N, E79K and E79R mutants, respectively, and a representative image is shown. **c**, Relative cell viability of E76K-expressing IMR-90 SV40-T cells compared to WT-expressing cells across a library of 336 epigenetic compounds. Cell viability was measured on day 4 post-treatment. Data represent the relative fold-change of the means between E76K and WT from three independent plates per condition. Similar enrichment for Aurora kinase and BET inhibitors was observed in two independent screening experiments. **d**, Venn diagram showing compounds that increased vulnerability in each mutant-versus WT-expressing cells. Bar plot highlights targets of compounds consistently reducing viability across four mutants. **e**,**g**, Dose–response curves for SNS-314 mesylate (Aurora kinase inhibitor) and CPI-203 (BET inhibitor) in IMR-90 SV40-T cells expressing H2B WT, mutants or empty vector, with corresponding IC_50_ values. Data represent means from three replicates, each consisting of six wells per condition. *P*-values were calculated using unpaired, two-tailed Student’s *t*-tests on IC_50_ values and on the means of all mutants compared to Empty and WT at the bottom plateau from nonlinear regression. **f**, Cell viability after three days of siRNA treatment targeting AURKA or AURKB in IMR-90 SV40-T cells expressing H2B WT, mutants or empty vector. Data represent mean ± s.d. from three replicates, each consisting of 14 wells per condition. *P*-values were calculated using unpaired, two-tailed Student’s *t*-tests comparing the means of all mutants to Empty and WT. **h**, Cell viability assay with 0.1 μM nocodazole in IMR-90 SV40-T cells expressing H2B WT, mutants or empty vector. Data represent mean ± s.d. from three replicates, each consisting of 16 wells per condition. *P*-values were calculated using unpaired, two-tailed Student’s *t*-tests comparing the means of all mutants to Empty and WT at the final time point. Data are representative of two independent experiments.

To more broadly assess stress sensitivity in human cells, we screened 336 compounds targeting epigenetic regulators. Mutant-expressing cells were generally more sensitive, showing pronounced vulnerability to Aurora kinase inhibitors and to bromodomain and extraterminal (BET) inhibitors (**Fig. 4c,d and Extended Data Fig. 8e**). For Aurora kinase inhibitors (SNS-314, Barasertib and GSK1070916), drug tolerance assays confirmed reduced tolerance, although IC_50_ values remained unchanged (**Fig. 4e and Extended Data Fig. 8f**). Knockdown of *AURKA* or *AURKB* phenocopied this reduced tolerance (**Fig. 4f**). By contrast, for BET inhibitors (CPI-203, OTX-015 and (+)-JQ-1), both tolerance and IC_50_ were reduced (**Fig. 4g and Extended Data Fig. 8g**), indicating heightened sensitivity to epigenetic perturbation. Both Aurora kinase and BET inhibition ultimately converge on the pathways controlling mitotic progression^49,50^. Indeed, mutant-expressing cells exhibited modest G2/M-phase and polyploid cell accumulation, which was further exacerbated under stress (**Extended Data Fig. 8h,i**), and showed decreased tolerance to mitotic inhibitors such as nocodazole and paclitaxel (**Fig. 4h and Extended Data Fig. 8j**). Together, these findings demonstrate that nucleosome instability compromises cellular resilience to environmental, epigenetic and mitotic stress, in part through reduced epigenomic robustness and increased mitotic vulnerability.

## Discussion

Our findings identify nucleosome stability as a core epigenetic safeguard that preserves cell identity, supports lineage specification, maintains stress resilience and sustains healthy tissue function throughout life. While nucleosome dynamics are regulated by histone variants, histone chaperones and chromatin remodelers and modifiers^51^, it has remained unclear how intrinsic nucleosome stability influences fundamental cell and tissue phenotypes that shape aging. We reveal that weakening of H2B–H4 interactions is sufficient to perturb lineage-specific gene expression programs and compromise cellular stress responses in a manner reminiscent of aging—without broadly affecting global chromatin accessibility, DNA damage, cell proliferation or viability. These results challenge and extend the conventional view that cell identity loss occurs downstream of cell cycle arrest, persistent DNA damage, global chromatin remodeling, senescence or niche alterations, which represent hallmark features of aging^1,7,52–55^. Together, our data reveal nucleosome stability itself as a key intrinsic mechanism safeguarding cell identity and lineage fidelity.

Nucleosome instability compromised some lineage-specific gene expression programs reminiscent of aging. Skeletal muscle displayed reduced fast-twitch myogenesis^56^; heart showed impaired fatty acid metabolism^57^; HSPCs exhibited a transcriptional bias toward megakaryocyte/erythroid lineages^31–34^; and human fibroblasts underwent a myofibroblast-like mesenchymal drift^46,58^. These tissue-specific alterations suggest that some lineage programs exist in “metastable” states maintained by nucleosome stability. Loss of this barrier increases fluctuations in these states, driving transcriptional drift and cell identity loss. Consistently, scRNA-seq analysis in the human fibroblast model revealed expansion of a cluster enriched for stress and myofibroblast-like identity programs, indicative of cell-state transition from metastable states. This framework aligns with previous studies linking chromatin dynamics to transcriptional noise^59^, cell fate metastability^60^ and age-associated transcriptional drift^61^. Here, we extend these findings by showing that intrinsic nucleosome stability safeguards cell identity and lineage fidelity, possibly by maintaining metastable lineage programs.

Nucleosome instability triggered intrinsic inflammatory and stress pathways, including TNF–NF-κB, interferon and p53 signaling, without broad genotoxicity or global changes in chromatin accessibility, indicating that these responses arise from decreased intrinsic nucleosome stability rather than canonical damage pathways, such as persistent DNA damage^55^. These findings highlight a previously unrecognized role for nucleosome stability in maintaining balanced cellular stress signaling. Furthermore, in human cells and yeast, nucleosome instability reduced resilience to exogenous challenges, encompassing environmental, epigenetic and mitotic stressors. Specifically, cells exhibited reduced tolerance to Aurora kinase inhibitors, linked to heightened mitotic vulnerability. Meanwhile, increased sensitivity to BET inhibitors may reflect an enhanced reliance on BET-regulated transcriptional programs for cell survival^62^. Collectively, these results indicate that nucleosome instability functions as an intrinsic form of stress, provoking epigenetic vulnerability and diminishing cellular resilience.

The impact of nucleosome instability extends from the cellular to the organismal level. In worms and flies, destabilizing nucleosomes shortened lifespan and accelerated the onset of age-associated transcriptomic and behavioral deficits. These observations suggest that nucleosome stability serves as an evolutionarily conserved safeguard, as its loss impairs cellular resilience and accelerates organismal aging.

Mutations at the H2B–H4 interface have been identified in various cancers^13–15^, yet their functional impact remains poorly understood. Given that both altered cell identity and stress signaling are hallmarks of cancer cells^63^, our data suggest that nucleosome instability may prime cells for malignant transformation by weakening tumor-suppressive differentiation programs and activating pro-tumorigenic inflammatory and stress signaling pathways. This framework unifies cell identity drift and stress signaling under a common epigenetic dysregulation that could predispose cells to cancer. Further studies are needed to elucidate the functional relationship between nucleosome instability and cancer development.

A key limitation of this study is that the impact of nucleosome instability varies across tissues, histone residues and model organisms. For instance, liver exhibited relatively modest effects compared to skeletal muscle and heart. In worms, the E73R mutant shortened lifespan, whereas the weaker E73K mutant had no measurable effect. Moreover, because histones are highly abundant proteins encoded by more than ten genes in human, mouse, fly and worm genomes, our experimental systems replaced less than 50% of endogenous histones with mutant variants. Therefore, both the quantitative extent of histone replacement and the severity of nucleosome instability likely contribute to the observed phenotypic variability.

Future work should define the molecular determinants of nucleosome stability and its homeostasis; examine how susceptibility varies across cell types, tissues and according to the degree of nucleosome destabilization; determine whether nucleosome perturbations act as primary drivers of aging; and explore whether stabilizing nucleosome conformations can counteract age- and cancer-associated phenotypes. Addressing these questions could further establish nucleosome integrity as a tractable target for interventions to extend healthspan and reduce disease risk.

## Methods

### Plasmid preparation

Point mutations in human histone H2B genes (coding region of *H2BC5* (NM_138720.2)) (WT, D68N, E76K, E76R, D68N/E76K) were cloned into pLenti CMV EGFP puro vector (addgene #17448) at BamHI-SalI site, which removed EGFP from the plasmid, with either C-terminal Flag tag. For EGFP or SNAP tag vectors, H2B or H2A (coding region of *H2AC4* (AK311785.1)) was cloned into pLenti PGK EGFP puro vector at BamHI-SalI site, where the CMV promoter was replaced by human PGK promoter from pLKO.1-blast (addgene #26655) at ClaI-BamHI site. EGFP/SNAP was fused C-terminally to H2B and N-terminally to H2A. Simian virus 40 large T antigen was cloned from pBABE SV40 zeocin T-antigen ER (kindly provided from Robert Weinberg) at BamHI site and inserted into pLenti PGK EGFP blast vector, where puromycin resistant gene was replaced by blasticidin resistant gene from pLKO.1-blast. For transduction of mouse HSPCs, mouse H2B genes (coding region of H2bc4 (NM_023422.3)) (WT, D68N, E76K, E76R) were cloned into MSCV PIG (Puro IRES GFP) (kindly provided by Aniruddha Deshpande, addgene #18751) at BglII-XhoI site with C-terminal Flag tag. For transduction of mouse tissues, mouse H2B (coding region of *H2bc4* (NM_023422.3)) (WT, D68N, E76K, E76R) were cloned into single-stranded AAV vector (VectorBuilder) with C-terminal Flag tag under the control of CMV promoter and Woodchuck hepatitis virus posttranscriptional regulatory element (WPRE). All constructs were verified by Sanger sequencing.

### Viral preparation

Lentiviruses were generated from 293 T cells transfected with expression vector, VSV-G envelope vector and psPAX2 packaging vector using lipofectamine 2000 (Invitrogen 52887). For retrovirus preparation, 293 T cells were transfected with expression vector and Ecopack gag/pol/env plasmid (a gift from Aniruddha Deshpande) using lipofectamine 2000. The virus supernatant was collected over 2 days, filtered through 0.45 μm membrane and directly used for the experiment.

Large-scale rAAV productions were carried out by the SBP Functional Genomics Core Facility. Briefly, HEK-293T cells were screened and optimized to establish high-titer, virus-producing clonal lines using Biomek i7 automation (Beckman Coulter). For production at scale, twenty 150 cm^2^ plates of low-passage, freshly cultured clonal 293T cells were co-transfected with rAAV vector, RepCap9 (a gift from James M. Wilson; addgene, 112865) and Helper (Cell Biolabs) plasmid DNA using polyethylenimine (Polysciences). At 84 h post-transfection, crude viral particles were harvested from the supernatant by polyethylene glycol precipitation and from the cell pellet through sonication, followed by benzonase (MilliporeSigma) treatment. Unconcentrated viruses were then purified by iodixanol (MilliporeSigma) gradient ultracentrifugation. The fraction containing concentrated rAAV particles was collected, buffer-exchanged into PBS, aliquoted and frozen at −80°C for long-term storage. Viral genome titers were measured via SYBR Green-based qPCR (Roche Diagnostics).

### Cell culture

IMR-90 SV40-T cells were maintained in Dulbecco’s modified Eagle’s medium (Gibco, 10313-121) supplemented with 10% fetal bovine serum (Corning, 35-011-CV), 2 mM L-glutamine, and 1% penicillin/streptomycin (Gibco, 15140-122). Cells were cultured at 37 °C in 5% CO_2_ and 5% O_2_. Stable cell lines expressing wild-type (WT) or mutant (D68N, E76K, E76R, D68N/E76K) were generated via lentiviral transduction followed by selection with puromycin (1 µg/ml) for two days.

For stress resilience assays, cells were either cultured continuously at 40 °C in 5% CO_2_ and 5% O_2_ or exposed once to ionizing radiation at a dose of 7 Gy (Rad Source RS-2000), followed by continued culture under the same conditions. For gene knockdown experiments, cells were transfected using Lipofectamine RNAiMAX (Thermo Fisher Scientific) with siRNAs targeting AURKA (siAURKA-1 (Invitrogen, s196); siAURKA-2 (Invitrogen, s197)) and AURKB (siAURKB-1 (Invitrogen, s17611); siAURKB-2 (Invitrogen, s17612)) or a non-targeting control (siCtr (Dharmacon, D-001810-01-05)) at a final concentration of 5 nM. Transfected cells were incubated for three days prior to viability assessment using CellTiter-Glo (Promega), with luminescence detected on an EnVision 2103 multilabel reader (PerkinElmer). All stress assays and siRNA treatments were performed in 384-well plates (Greiner Bio-One 781098). Cells and reagents were dispensed using a Multidrop Combi reagent dispenser (Thermo Fisher Scientific).

As a positive control for double-stranded DNA breaks, cells were treated with etoposide at a final concentration of 10 μM for 24 h prior to fixation.

### Drug tolerance assay

Cells were seeded into 384-well plates and treated with increasing concentrations of Aurora kinase inhibitors (SNS-314 Mesylate (SelleckChem, S8699); Barasertib (SelleckChem, S1147); GSK1070916 (MedChemExpress, HY-70044)) and BET inhibitors (CPI-203 (Sigma, SML1212); OTX-015 (Sigma, SML1605); (+)-JQ-1 (MedChemExpress, HY-13030)). Cells were treated for three days for Aurora kinase inhibitors and four days for BET inhibitors prior to viability assays. Paclitaxel (Cayman Chemical, 10461) and nocodazole (MedChemExpress, HY-13520) were also used in selected assays. After drug treatment, cell viability was measured using CellTiter-Glo, and luminescence was detected with an EnVision 2103 multilabel reader. Drug sensitivity was assessed using GraphPad Prism by calculating IC_50_ values from nonlinear regression of cell viability curves, while drug tolerance was quantified as the fitted bottom asymptote, representing residual viability at high drug concentrations. Sensitivity to nocodazole and paclitaxel was evaluated over a multi-day time course by monitoring luminescence-based viability.

### Epigenetic library screen

A high-throughput screen of an epigenetic drug library was performed to identify compounds that selectively impaired viability in mutant cell lines. The library was provided by the SBP Conrad Prebys Center for Chemical Genomics. Compounds from the Epigenetics compound library and DMSO controls were transferred into 384-well plates using an Echo 655 Liquid Handler (Beckman Coulter). The drug concentration was 1 μM (2.5 nL of 10 mM drugs). Cells were incubated for four days, and viability was measured using CellTiter-Glo with an EnVision 2103 multilabel reader. Relative viability was calculated following drug treatment and plotted against statistical significance (−log₁₀ P-value). Comparative analysis across mutants was visualized using Venn diagrams and bar graphs to identify overlapping drug sensitivities.

### Cell cycle assay

Cell cycle distribution was assessed using the Click-iT Plus EdU Pacific Blue Flow Cytometry Assay Kit (Invitrogen, C10636), following the manufacturer’s instructions. Cells were subjected to either heat stress (40 °C for 24 h) or irradiation (7 Gy) 24 h prior to the experiment. EdU incorporation was used to label cells undergoing DNA synthesis, and total DNA content was counterstained with propidium iodide (PI; Invitrogen, F10797). Flow cytometry was performed on a Novocyte flow cytometer (ACEA Biosciences Inc.), and data were analyzed using FlowJo software. Cell cycle phases (G1, S, G2/M) were determined by EdU incorporation and PI intensity. Populations with DNA content greater than 4N were also quantified to assess potential re-replication or polyploidy.

### Sequential salt fractionation

Sequential salt fractionation was performed, as previously described^64^. Briefly, 3 × 10^6^ IMR-90 SV40-T cells expressing WT or mutant H2B were lysed in 100 μl of modified buffer A (25 mM HEPES pH 7.6, 25 mM KCl, 5 mM MgCl_2_, 0.05 mM EDTA, 0.1% NP-40, 10% glycerol) on ice. Lysates were centrifuged, and the supernatant was collected as the cytoplasmic fraction. Pellets were washed once with 0 mM NaCl mRIPA (50 mM Tris-HCl pH 8.0, 1% NP-40, 0.25% sodium deoxycholate), then sequentially resuspended in mRIPA containing 0, 0.6, 0.9 or 1.2 M NaCl and centrifuged at 12,000 × *g* for 5 min. Nuclear extracts were collected at each step. Samples were resuspended in Laemmli sample buffer and used for western blotting.

### Fluorescence recovery after photobleaching (FRAP)

IMR-90 SV40-T cells expressing H2B-EGFP constructs were seeded on PhenoPlate^TM^ 96-well microplates (PerkinElmer) one day prior to the experiment. EGFP fluorescence was imaged using a Nikon N-SIM E super-resolution/A1 ER confocal microscope with a 60 × oil immersion objective under temperature control (37 °C). A rectangular region of interest, approximately half of the nucleus, was bleached with the 488-nm laser at full power for 45 s, and recovery was monitored at 4 min intervals for 60 min. Fluorescence intensities at the bleached site were corrected using the unbleached region of the same nucleus. Recovery curves were fitted with a single-exponential model in GraphPad Prism to derive plateau values (*Ymax*). More than fifteen nuclei were analyzed per condition.

### Histone turnover assay

IMR-90 SV40-T cells expressing H2B-SNAP constructs were labeled with 1.5 μM SNAP-Cell TMR-Star (New England Biolabs #S9105) in DMEM for 30 min in the incubator, washed with PBS, and blocked with 5 μM SNAP-Cell Block (New England Biolabs #S9106) in DMEM. After a 0–30 h chase period, residual nuclear fluorescence was imaged on a Nikon T2 microscope with automated image capture. Cells were then fixed and counterstained with DAPI. Images were analyzed in NIS Elements AR v5.21.03 using dark background subtraction, thresholding, size exclusion and automated partitioning to identify nuclear DAPI features. At least 2,600 nuclei were analyzed per condition. Signal intensities were normalized to time 0 and fitted to a one-phase exponential decay model in GraphPad Prism. The degradation rate constant (*k*) was obtained from the fit, and the protein half-life (*t*_1/2_) was calculated as ln2/*k*. To ensure accurate modeling of the decay, the plateau parameter was constrained to zero during curve fitting.

### Mouse models

Male 3-week-old C57BL/6J mice (The Jackson Laboratory, strain #000664) were used. A total of 100 μl of rAAVs (1-5 × 10^12^ VG) were administered via retro-orbital injection by Sanford Burnham Prebys Medical Discovery Institute Animal Facility. Mice were housed five per cage and maintained under controlled temperature (22.5 °C) and a 12 h light/dark cycle, with food and water provided *ad libitum*. Four weeks post-transduction, mice were euthanized with CO_2_, and then quadriceps skeletal muscle, heart and liver were collected, snap-frozen in liquid nitrogen and stored at –80 °C. All animal procedures were approved by the Institutional Animal Care and Use Committee (IACUC) of Sanford Burnham Prebys Medical Discovery Institute, and all experiments were performed at Sanford Burnham Prebys Medical Discovery Institute Animal Facility in compliance with the IACUC guidelines and relevant ethical regulations for animal research.

### Histology and H&E staining

Tissues were paraffin-embedded, sectioned at 5 µm, and mounted on glass slides. Slides were dried at 58 °C for 1 h, deparaffinized, and rehydrated through graded alcohols. Hematoxylin and eosin (H&E) staining was performed using a Leica Autostainer XL (Leica Microsystems) following standard protocols. All histological processing and staining were carried out by the Histology Core Facility at Sanford Burnham Prebys Medical Discovery Institute.

### HSPCs isolation from mouse bone marrow

Bone marrow cells were isolated from spinal cord and hind leg bones of 6 male, 9-week-old C57BL/6 mice. Collected bones were placed in 15% FBS/PBS, then crushed using a pestle in a mortar and filtered through a 40 μm cell strainer (Fisher Scientific). Cells were pelleted by centrifugation, resuspended in lysis buffer (BD Biosciences, 555899) and incubated for 4 min to lyse red blood cells. After washing twice with 15% FBS/PBS, lineage-positive cells were depleted using the EasySep Mouse Hematopoietic Progenitor Cell Isolation Kit (STEMCELL, #19856). Briefly, mouse FcR blocker was added to the sample, followed by the hematopoietic progenitor cell isolation cocktail and incubated for 15 min at 4 °C. Streptavidin RapidSpheres were then added and incubated for 10 min before placing the sample in the magnet; the supernatant containing lineage-negative cells was collected. Cells were pelleted, resuspended at 20 × 10^6^ cells/ml in PBS containing 2% FBS and 1 mM EDTA, and subjected to flow cytometric sorting.

Sorting was performed using a FACSAria III cell sorter (BD Biosciences, P0024) with FACSDiva software v8.0.1. Biotin-positive cells were excluded by gating with a BV421 fluorescence minus one (FMO) control [405/450 50-A], and subsequent gates were set using PE and APC FMOs to define Sca-1⁺ and c-Kit⁺ populations, respectively. FMOs were used to establish gating thresholds and discriminate true positive signals from background in populations with heterogeneous or dim marker expression. Lin⁻Sca-1⁺c-Kit⁺ (LSK) cells were collected, yielding 29,510 cells from a total of 35,115,973 bone marrow cells. Antibodies used were PE anti-mouse Ly-6A/E (Sca-1) (rat monoclonal antibody, BioLegend, #108107), APC anti-mouse CD117 (c-Kit) (rat monoclonal antibody, BioLegend, #105811), and Brilliant Violet 421 Streptavidin (BioLegend). For fluorescence compensation, single-stained controls for PE, APC, and BV421, along with an unstained control, were prepared prior to sample acquisition and used to generate a compensation matrix.

### Myeloid and megakaryocyte/erythroid lineage differentiation

HSPCs were cultured in DMEM supplemented with 15% FBS, 2 mM L-glutamine, and 1% penicillin/streptomycin, 20 ng/ml mouse SCF (Thermo Fisher Scientific, 250-03), 10 ng/ml mouse IL-6 (Thermo Fisher Scientific, 216-16) and 6 ng/ml mouse IL-3 (Thermo Fisher Scientific, 213-13). One day post-isolation, cells were transduced with MSCV retroviruses for 1 day on a retronectin-coated plate, followed by puromycin selection (2.5 μg/ml) for additional 2 days. Colony-forming assays were performed using MethoCult GF M3434 (STEMCELL Technologies), according to manufacturer’s instructions. Briefly, approximately 5,500 cells were mixed with 4 ml of methylcellulose media supplemented with 20 ng/ml mouse FLT3L (Thermo Fisher Scientific, 250-31L) and 50 ng/ml mouse TPO (Thermo Fisher Scientific, 315-14), and plated in triplicate (1.1 mL per 3.5 mm dish) using a 16-gauge blunt-end needle (STEMCELL Technologies, 28110). Colonies were manually counted under bright-field microscopy and classified as CFU-G, CFU-M, or CFU-GM based on morphology. Cells were harvested from methylcellulose cultures 9 days after differentiation using DPBS and stored at −80 °C for mRNA-seq or used for flow cytometry. To assess differentiation efficiency, cells were stained with antibodies against lineage markers, Sca-1 (BioLegend, 108119), c-Kit (BioLegend, 105811), Gr-1 (BioLegend, 108433) and Mac-1 (BioLegend, 101212). Flow cytometry was performed, and data were using a Novocyte (ACEA Biosciences Inc.), analyzed with FlowJo software. Quantification of Sca-1⁺c-Kit⁺, Gr-1⁺ and Mac-1⁺ populations was used to assess differentiation efficiency.

### Western blotting

For protein expression analysis, cell suspensions were directly lysed in 2 × Laemmli sample buffer (125 mM Tris-HCl pH 6.8, 4% SDS, 20% glycerol, 0.2% bromophenol blue). Mouse tissues and Drosophila samples were homogenized using tissue grinding tubes (Precellys, P000917-LYSK0-A for skeletal muscle and heart; P000912-LYSK0-A for liver and flies) with a Precellys Evolution homogenizer (Bertin Technologies) for three cycles of 4 × 30 s at 6,000 × *g*. Lysates were centrifuged at 12,000 rpm for 5 min, and the supernatant was boiled at 95 °C for up to 5 min and stored at −80 °C. Protein samples (5–50 μg per lane) were separated by SDS–PAGE using 15% Criterion Tris-HCl Protein Gels (Bio-Rad, 3450020) and Tris/Glycine/SDS buffer (Bio-Rad, 1610772) and transferred to nitrocellulose membranes (Bio-Rad, 1620112) using the Trans-Blot Turbo system (Bio-Rad). Membranes were block with 5% skim milk and incubated with primary antibodies overnight at 4 °C, followed by HRP-conjugated secondary antibodies for 1 h at room temperature. Bands were visualized using enhanced chemiluminescence reagents (Thermo Fisher Scientific, 34580 or 34095) and imaged with the ChemiDoc Imaging System (Bio-Rad).

### Immunofluorescence

Cells were seeded on PhenoPlate 96-well microplates (PerkinElmer) and fixed with 10% neutral buffered formalin (Epredia, 9400-1). Fixed cells were permeabilized with buffer (0.2% Triton X-100 and 0.5% BSA in PBS) for 10 min, blocked with 0.5% BSA in PBS and incubated with primary antibodies diluted in 0.2% BSA in PBS for 1 h at room temperature. After two washes with 0.2% BSA in PBS, samples were incubated with secondary antibodies for 1 h. Images were acquired using a Nikon T2 microscope with manual or automated image capture. Image analysis was performed in NIS Elements AR v5.21.03 using dark background subtraction, thresholding, size exclusion and automated partitioning to identify nuclear DAPI features. Images were processed using Fiji^65^.

### Antibodies

For western blotting, the following primary antibodies were used: Flag M2 (Sigma, F3165, 1:5000; Cell Signaling, 14793, 1:5000), H2B (Cell signaling, 12364, 1:5000), GFP (Abcam, 6556, 1:1000) and GAPDH (Santa Cruz, sc-47724, 1:10000; Abcam, 9485, 1:10000). For immunofluorescence, the following primary antibodies were used: Flag M2 (Sigma, F3165, 1:1000; Cell Signaling, 14793, 1:1000), Phospho-Histone H2A.X (Ser139) (Millipore, 05-636, 1:1000) and Histone H2A.X (Abcam, ab11175, 1:1000). The following secondary antibodies were used: Goat anti-Mouse IgG, IgM (H+L) HRP (Thermo Fisher Scientific, 31446), Goat anti-Rabbit IgG, (H+L) HRP (Millipore, AP307P), Goat anti-Mouse IgG (H+L), Alexa Fluor 488 (Thermo Fisher Scientific, A11029), Goat anti-Rabbit IgG (H+L) and Alexa Fluor 594 (Thermo Fisher Scientific, A11012).

### RNA extraction, cDNA synthesis and qPCR

Total RNA from human IMR-90 SV40-T cells and mouse HSPCs was extracted using the Quick-RNA Miniprep Kit (Zymo Research, R1055), according to the manufacturer’s instructions. Eluted RNA was quantified using a Nanodrop 2000 spectrophotometer (Thermo Fisher Scientific) and reverse-transcribed into cDNA using the SuperScript III First-Strand Synthesis System (Invitrogen, 18080093) following the manufacturer’s instructions. qPCR was performed using SYBR green master mix (Applied Biosystems, A25742) in 384-well plates. Cycling conditions were 95 °C for 20 s, followed by 40 cycles of 95 °C for 1 s and 60 °C for 20 s, with a melt-curve analysis. The following primers were used: H2B-F (ACCAAGGCCGTCACCAAGTAC) and Flag-R (CTTGTCGTCATCGTCTTTGTAGTCTC).

For mouse tissues, frozen samples (≤0.5 × 0.5 × 0.5 mm) were placed in tissue grinding tubes (Precellys, P000917-LYSK0-A for skeletal muscle and heart; P000912-LYSK0-A for liver) containing TRIzol Reagent (Invitrogen, 15596026) and homogenized in a Precellys Evolution homogenizer (Bertin Technologies) for three cycles of 4 × 30 s at 6,000 *g*. Lysates were centrifuged at 12,000 rpm for 1 min, and the supernatant was collected. RNA was then extracted using the Direct-zol RNA Miniprep Kit (Zymo Research, R2052), according to the manufacturer’s instructions.

### RNA-seq and data analysis

For mouse and worm samples, PolyA RNA isolation and library preparation was performed with the Watchmaker mRNA Library Prep Kit (Watchmaker Genomics: 7BK0001) with xGEN Stubby adaptors (IDT, 336338420) and xGEN 10nt UDIs (IDT, 336338436). Libraries were sequenced with the Element Biosciences AVITI Sequencing platform using the AVITI 2×75 High Output Cloudbreak Freestyle Kit (Element Biosciences, 860-00015). For human IMR-90 SV40-T cell samples, PolyA RNA isolation and library preparation was performed with NEBNext Ultra II DNA Library Prep Kit for Illumina (New England Biolabs, E7645) with NEBNext Multiplex Oligos for Illumina (96 Index Primers) (New England Biolabs, E6609). Libraries were sequenced with Novaseq6000 at PE100 at the UC San Diego IGM Genomics Center.

Reads were trimmed for adaptors and low-quality bases (Q<20) using Trimmomatic (Galaxy v0.38.0) or Trim Galore! (Galaxy v0.6.7+galaxy0 or v0.6.7+galaxy1) and then aligned to the human (hg38), mouse (mm10) or worm (ce10) reference genome using HISAT2 (Galaxy v2.2.1+galaxy1). Aligned reads were quantified with featureCounts (Galaxy v2.0.3+galaxy1 or v2.0.3+galaxy2) using exon regions from gene annotation files (human: GENECODE v32; mouse: UCSC mm10; worm: NCBI Refseq ce11). Differential expression analysis was performed with DESeq2 (Galaxy v2.11.40.7+galaxy2 or v2.11.40.8+galaxy0) using the local fitting strategy.

Principal component analysis (PCA) was performed in R (v4.4.2) using the prcomp function on log_2_-transformed, normalized count data. Genes with zero variance across samples were excluded prior to PCA. The first two principal components (PC1 and PC2) were visualized using the ggplot2 package (v3.5.2), with genotype-based color coding and axes indicating the proportion of variance explained. Hierarchical clustering of differentially expressed genes (DEGs) was performed using Euclidean distance and complete linkage via the pheatmap package (v1.0.12), with rows (genes) clustered, columns (samples) fixed and *Z*-scores capped to limit extreme values. Pathway enrichment analysis was performed for hallmark and cell type signature gene sets, including Tabula Muris Senis gene sets, using GSEA 4.4.0 (Broad Institute). Upstream regulator analysis was performed using Ingenuity Pathway Analysis (IPA; QIAGEN). GSEA results were visualized as bubble plots: for hallmark signatures, point size represented –log10(FDR) and color indicated normalized enrichment score (NES); for cell type signatures, NES was plotted on the x-axis, cell type on the y-axis, point size represented –log10(FDR) and color indicated gene set size. For heatmap analysis, gene expression values were log_2_-transformed and standardized to *Z*-scores on a per-gene basis, and visualized using the ComplexHeatmap package (v2.22.0) with color scaling from circlize (v0.4.16), clustering of rows (genes) and a fixed column (samples) order. To compare gene groups, *Z*-scores of individual genes were averaged per sample to obtain mean *Z*-scores. Pearson correlation coefficients were calculated between these mean *Z*-scores and corresponding protein turnover values. Data preprocessing and manipulation were performed using dplyr (v1.1.4) and readr (v2.1.5). For immune signature analysis, bulk RNA-seq data were processed using CIBERSORTx with the LM22 signature matrix. Immune cell proportions were compared between control samples (Empty and WT) and mutant samples (D68N, E76K, and E76R) using Wilcoxon rank-sum tests. Results were visualized using boxplots with overlaid jittered points for each cell type.

### RNA-seq in primary human dermal fibroblasts

Primary human dermal fibroblasts from four young donors (age 23–33) and five aged donors (age 64–72) were obtained from the San Diego Nathan Shock Center (SD-NSC) Human Cell Models of Aging Core (see **Supplementary Table 1**). Cells were cultured in DMEM (Gibco, 11965092) supplemented with 10% fetal bovine serum (Corning, 45000-736), 1% non-essential amino acids (Gibco, 11140050) and 1% GlutaMAX (Gibco, 35050061) at 37 °C, 5% CO_2_ and ambient O_2_. For RNA-seq, 200,000 cells per cell line were seeded in single wells of 6-well plates and cultured for 48 h. RNA was extracted using the RNeasy Mini Kit (Qiagen, 74104), according to the manufacturer’s instructions. Briefly, cells were washed with Dulbecco’s phosphate-buffered saline (DPBS; Gibco, 14190094) and lysed in 350 μL RNeasy lysis buffer containing β-mercaptoethanol. Lysates were homogenized using QIAshredder columns (Qiagen, 79656), followed by purification with RNeasy Mini spin columns, including on-column DNase digestion. PolyA library prep was performed by the Salk Institute for Biological Studies Next Generation Sequencing and Genomics Core, and sequencing was carried out on an Illumina NovaSeq 6000 SP platform. Raw FASTQ files were mapped to hg19 (https://www.ncbi.nlm.nih.gov/grc/human) by STAR (v2.5.3a), and DEG analysis was performed with HOMER pipeline (v4.11.1).

### Single-cell RNA sequencing

IMR-90 fibroblasts expressing WT or mutant H2B were processed using the Chromium platform at the Center for Epigenomics at UC San Diego. Data were processed with the Cell Ranger pipeline (v8.0.0, 10x Genomics) for demultiplexing, alignment to the GRCh38 reference genome and generation of gene–barcode count matrices. Expression matrices were analyzed using Scanpy (v1.9.8). Cells with fewer than 200 detected genes, more than 5% mitochondrial gene expression, or more than 8,000 detected genes were excluded. Genes detected in fewer than three cells were also removed. Mitochondrial and ribosomal genes were excluded due to their disproportionate expression. The filtered data were normalized to a total of 1 × 10^6^ transcripts per cell and log-transformed to stabilize variance. Highly variable genes were selected (minimum mean = 0.5, maximum mean = 8, minimum dispersion = 0.5). Principal component analysis (PCA) was applied, and the top 40 principal components were used for neighborhood graph construction and clustering with the Leiden algorithm (resolution = 1). Uniform Manifold Approximation and Projection (UMAP) was used for visualization in two dimensions. Cluster marker gene identification and differential expression analysis between WT and E76K or E76R mutants were performed using the Wilcoxon rank-sum test.

### MNase digestion

Global chromatin accessibility was assessed with micrococcal nuclease (MNase) digestion. A total of 1 × 10^6^ freshly harvested cells were resuspended with modified RIPA buffer (50 mM Tris-HCl pH 8.0, 1% NP-40, 0.25% sodium deoxycholate), and the pellet was resuspended with 4 Kunitz U/100 μl MNase (NEB, M0247) and incubated at 37 °C for 10–30 min. DNA was extracted using the Quick-DNA Miniprep Kit (Zymo Research, D3024) and analyzed by gel electrophoresis on a 0.8% agarose gel.

### ATAC-seq

ATAC-seq was performed in IMR-90 SV40-T cells expressing WT or E76K using ATAC-seq Kit (Active Motif, 53150). For TGFβ treatment, cells were treated with 2 ng/ml TGFβ1 24 h before the experiment. Briefly, 1 × 10^5^ cells were washed with PBS, resuspended in ATAC-seq lysis buffer and incubated with the Tagmentation master mix at 37 °C for 30 min in a thermomixer set at 1,000 rpm. Tagmended DNA was purified, and libraries were prepared with i5 and i7 indexes. Size selection was performed using AMPure XP beads (Beckman Coulter, A63881) with a 0.5–1.3× ratio. Libraries were quantified with the KAPA library quantification kit (Roche) and sequenced on a NovaSeq 6000 (Illumina) at the UC San Diego IGM Genomics Center.

Raw sequencing reads were assessed for quality using FastQC (v0.10.0). Adapter sequences and low-quality bases were trimmed with Trim Galore (v0.3.0), and the resulting clean reads were aligned to the GRCh38 reference genome using Bowtie2 (v2.4.2). Aligned reads were sorted and indexed with SAMtools (v1.6), and PCR duplicates were identified and removed using the MarkDuplicates function in Picard Tools (v2.27.5). Peaks were called using MACS2 (Galaxy v2.2.7.1+galaxy0) from all conditions with or without TGFβ treatment, and differential peak detection was performed using featureCounts (Galaxy v2.0.1+galaxy2) followed by DESeq2 (Galaxy v2.11.40.8+galaxy0) across 114,678 peak regions. Gene annotation was performed using GREAT^66^ (v4.0.4), assigning each peak to the nearest genes within ± 5 kb or up to 100 kb. Plot profiles were generated using deepTools computeMatrix (Galaxy v3.5.4+galaxy0) followed by plotProfile (Galaxy v3.5.4+galaxy0).

### SLAM-seq

SLAM-seq^67^ was performed in IMR-90 SV40-T cells expressing WT or mutant H2B using the SLAMseq Kinetics Kit – Anabolic Kinetics Module (Lexogen, 061.24), according to the manufacturer’s protocol. A total of 2 × 10^5^ cells were seeded in a well of a 6-well plate, and media was changed the following day. After an additional day, cells were treated with 100 μM 4-thiouridine (4sU) for 6 h to label newly synthesized RNA. Total RNA was extracted using the Quick-RNA Miniprep Kit (Zymo Research, R1055), followed by alkylation with iodoacetamide (IAA) to induce T>C conversions. RNA libraries were prepared using the QuantSeq 3′ mRNA-Seq V2 FWD Library Generation Module with Lexogen UDI 12 nt Set A1 and amplified using the Library Amplification Module. Libraries were quantified with the KAPA Library Quantification Kit (Roche) and sequenced on a NovaSeq 6000 (Illumina) at the UC San Diego IGM Genomics Center.

Reads were trimmed for low-quality bases (Q<20) using Trimmomatic (Galaxy v0.38.0) and aligned to the GRCh38 reference genome. T>C conversions were quantified using the SLAM-DUNK pipeline (Galaxy v0.4.3+galaxy1), which includes filtering, mapping, mutation calling and quantification modules. Differential expression analysis was performed using DESeq2 (Galaxy v2.11.40.8+galaxy0). Transcripts with significant changes in T>C conversion rates were considered transcriptionally regulated.

### Worm strain construction and maintenance

CRISPR–Cas9 genome editing was used to introduce the E73K and E73R mutations into the endogenous *his-41* locus of the N2 Bristol background. Strains PHX8148, PHX8245 and PHX8202 were generated by Suny Biotech. PHX8148 is a wild-type strain in which the *his-41* E73 codon was replaced with E73E (PHX8148: AATTGCTTCGGAA). The E73K mutation was introduced into PHX8245 (*his-41*(syb8245)); PHX8245: AATTGCTTCGAAA), and the E73R mutation was introduced into PHX8202 (*his-41*(syb8202)); PHX8202: AATTGCTTCGAGA). Worms were maintained on nematode growth medium (NGM) agar plates seeded with *Escherichia coli* strain OP50 at 20 °C. The N2 Bristol strain was obtained from the Caenorhabditis Genetics Center (CGC).

### Lifespan assay in worms

Lifespan assays were conducted following the protocol previously detailed^68^. In brief, L1 age-synchronized worms were plated into 96-well microtiter plates at a volume of 120 μl per well. At the L4 larval stage, animals were sterilized by adding FUDR to each well to a final concentration of 0.12 mM in a total volume of 150 μL. Worm populations were assessed for viability every 2–3 days after shaking plates for 5 min on a microtiter plate shaker to stimulate movement. Plates were stored in incubators maintained at 20 °C. Lifespan analysis was performed using STATA.

### RNA extraction from worms and RNA-seq data analysis

Worm populations were cultured in 15 cm dishes (three dishes per strain and time point), each containing 20 mL of S-complete medium. Worms were maintained at a density of 100 worms/ml and fed OP50 bacteria (ampicillin/carbenicillin-resistant) at a concentration of 4 mg/ml. To prevent contamination, Fungizone (0.1 μg/ml) and carbenicillin (50 μg/ml) were added to the medium. At the L4 stage, FUDR was added to sterilize the animals at a final concentration of 0.12 mM. Worms were harvested at the indicated time points, suspended in minimal PBS and stored at −80 °C overnight or until extraction. For lysis, worms were disrupted in TRIzol (Invitrogen, 15596-026) with zirconium and glass beads using a tissue homogenizer. RNA was extracted by phenol-chloroform separation and further purified using the RNeasy Plus Micro Kit (Qiagen, 74034).

To assess accelerated aging signatures, age-associated genes were first defined by differential expression between WT day 3 and day 10 adults. Genes upregulated with age in WT and already elevated in E73R mutants at day 3 were classified as accelerated age-up genes (n = 645). Conversely, genes downregulated with age in WT and already reduced in E73R mutants at day 3 were defined as accelerated age-down genes (n = 1089). Mean expression levels of these gene sets were calculated per condition, with 95% confidence intervals estimated as mean ± 1.96 × standard error. Gene annotation and functional enrichment analysis of the accelerated age-up and age-down gene sets were performed using Metascape^69^.

### Generation and maintenance of transgenic flies

Transgenes comprising 5×UAS/mini_Hsp70/H2B (His2B:CG17949, NM_165381.4; WT or E73K) C-terminally fused to a Flag tag were generated by VectorBuilder in an attB-containing plasmid backbone and integrated into genomic attP landing sites in *Drosophila melanogaster* via ΦC31 integrase (BestGene Inc.). Ubiquitous expression was achieved by crossing to Act5C-GAL4 drivers (Bloomington Drosophila Stock Center, stock #25374). Flies were maintained on a standard yeast, corn starch, and molasses diet (10% yeast, 12% sugar, 1.5% agar) at 25 °C under a 12 h light/dark cycle and controlled humidity.

### Fly lifespan and locomotor performance assays

Lifespan assays were performed, as previously described^70^. Between 140 and 365 flies per sex and genotype were housed at 25 flies per vial. Flies were transferred to fresh food every 2–3 days, and survival or censoring was recorded at each transfer. Kaplan–Meier survival curves were generated in GraphPad Prism, and significance was assessed by log-rank test.

Locomotor activity was assayed by negative geotaxis at three time points during lifespan, as previously described^71^. Groups of 3–27 flies were tapped to the bottom of the vial, and the fraction ascending above 2 cm within 10 s was scored. Each cohort was tested in triplicate, and four independent cohorts per sex and genotype were analyzed.

### Yeast strain construction and maintenance

Wildtype or mutant (*htb1* D71N, E79K or E79R) copies of the *HTA1*-*HTB1* gene cassette were integrated into the *HTA1*-*HTB1* locus of the *Saccharomyces cerevisiae* histone shuffle strain FY406 (MATa (*hta1*-*htb1*)Δ::LEU2 (*hta2*-*htb2*)Δ::TRP1 *lys2*-128 *leu2*Δ1 *ura3*-52 *trp1*Δ63 *his3*Δ200 pSAB6[URA3, CEN, ARS, *HTA1*-*HTB1*]) using standard transformation protocols. Transformants were selected on 5-fluoroortic acid, genotyped, and the sequence of *htb1* was confirmed by Sanger sequencing. Yeast strains were cultured in YPD medium at 30 °C, and growth was monitored by measuring optical density at 600 nm (OD_600_) at regular intervals to assess proliferation rates.

### Replicative lifespan and stress tolerance assays in yeast

Replicative lifespan was measured using a microfluidics-based platform adapted from the HYAA-Chip protocol^72^. Briefly, filter-sterilized YPD medium was loaded into an AD-Chip at 20 μL/min using 10 mL syringes (BD Biosciences) driven by a KDS-230 pump (KD Scientific) and maintained at 1 μL/min. Yeast cells were grown to mid-log phase, diluted 1:20 and manually loading into the AD-Chip. Multiposition time-lapse imaging was performed using an EVOS FL Auto system equipped with a 30 °C environmental chamber (Thermo Fisher), a 20 × objective and transmitted light optics. Images (three per channel) were acquired every 15 min for 65 h. Image series were analyzed using ImageJ (National Institutes of Health), and cell divisions were manually counted for at least 50 mother cells per strain. Replicative lifespan was defined as the mean number of daughter cells produced. Lifespan differences were assessed using a two-sided Wilcoxon rank-sum test.

Stress tolerance assays were performed using stationary-phase cells, as growth phase can influence stress sensitivity. Cells were plated onto YPD agar or YPD supplemented with stress-inducing agents and incubated under the control condition at 30 °C for 2 days or under stress conditions: heat shock at 42 °C for 16 h or UV irradiation (300 J/m²), followed by incubation at 30 °C until 2 days; DNA-damaging agents, including camptothecin (CPT) and methyl methanesulfonate (MMS) for 2 days; and low temperature at 15 °C for 5 days. Growth and survival were assessed by spotting 1:10 serial dilutions starting from 2 OD_600_ and imaging plates after incubation.

### Statistical analysis

No statistical method was used to predetermine sample size. Sample sizes for cell culture experiments were determined empirically for each experiment. Sample sizes for animal experiments were based on pilot studies and were similar to those used previously in comparable worm and fly experiments. Statistical analyses were performed using GraphPad Prism (v9), Microsoft Excel, STATA and R- or Python-based tools, except for sequencing-based analyses. Quantitative data are displayed as mean ± s.d. and represented as error bars unless otherwise indicated. Results from each group were averaged and used to calculate descriptive statistics. *P*-values < 0.05 were considered statistically significant. For cell culture and animal experiments, pairwise comparisons were performed using two-tailed Student’s *t*-tests, assuming equal variance between groups (Fig. 1b,c; 4a,e–h; Extended Data Fig. 1e,i; 2b; 4c,d; 8f,g,i,j). Survival analyses were performed using log-rank (Mantel–Cox) tests, and only adjusted *P*-values are reported (Fig. 3b,k; Extended Data Fig. 7j; 8d). For *C. elegans* mean lifespan comparisons, one-way ANOVA was used to assess group differences, followed by Tukey’s HSD-adjusted emmeans for pairwise comparisons (Fig. 3b). For locomotor performance assay, data were analyzed using two-way ANOVA (genotype × age) separately for each sex, followed by post hoc pairwise comparisons between the Control/WT group and mutant at individual ages using emmeans with Tukey’s HSD adjustment (Fig. 3l). Sequencing data were analyzed using standard pipelines. Differential gene expression was assessed using DESeq2, with genes showing FDR < 0.05 considered differentially expressed for mRNA-seq and differentially accessible for ATAC-seq (Fig. 3d; Extended Data Fig. 3a; 6c). For SLAM-seq, genes with FDR < 0.1 were considered differentially expressed (Extended Data Fig. 6g). GSEA was performed using MSigDB Hallmark and cell type signature databases, including Tabula Muris Senis aging signatures, with Desktop GSEA (v4.4.0). IPA (Qiagen) used a right-tailed Fisher’s exact test to determine statistical significance of upstream regulator enrichment. GSEA FDRs and IPA *Z*-scores are shown without a statistical cut-off to assess data trends unless otherwise indicated. Expression *Z*-scores were calculated using log2-transformed expression values followed by sample-wise *Z*-score normalization (mean = 0, SD = 1) (Fig. 2e; Extended Data Fig. 5b). Differential expression was assessed using log2 fold change and FDR (Extended Data Fig. 5f). To assess the relationship between chromatin dynamics and gene expression (Fig. 2f), Pearson correlation coefficients and linear regression lines were computed using GraphPad Prism. For scRNA-seq data, clustering and dimensionality reduction were performed using Seurat, and differential expression was assessed using Wilcoxon rank-sum tests with an FDR cut-off of 0.05 for cluster comparisons (Extended Data Fig. 5d). Estimated immune cell fractions from CIBERSORTx were compared between control and mutant groups using the Wilcoxon rank-sum test (unpaired, two-tailed) for each cell type. Investigators were double-blinded when measuring colony numbers in HSPC differentiation assays (Extended Data Fig. 4c). Otherwise, investigators were not blinded during cell culture experiments, animal experiments or outcome assessment.

## Data availability

All sequencing data have been deposited in GEO under accession codes: GSE307377 (mRNA-seq of human donors), GSE307414 (mRNA-seq of HSPCs), GSE307475 (mRNA-seq of IMR-90 SV40-T), GSE307833 (mRNA-seq of *C. elegans*), GSE307869 (ATAC-seq of IMR-90 SV40-T), GSE308172 (SLAM-seq of IMR-90 SV40-T), GSE308173 (scRNA-seq of IMR-90 SV40-T) and GSE308174 (mRNA-seq of mouse tissues). All other raw data can be obtained by contacting the corresponding authors.

## Author contributions

Conceptualization: H.T., P.D.A. Methodology: H.T., B.S.M, C.G., S.L., T.M., Z.M.C, A.L., R.A., A.R., M.G.T., L.H., A.D., A.D. Data generation and analysis: H.T., B.S.M, C.G., X.L., S.L., T.M.M., K.G., A.A., C.K., M.P. Writing: H.T., P.D.A. Overall supervision: H.T., P.D.A.

## Acknowledgement

This research was supported by grants R01 AG071464, P01 AG031862 and R01 AG071861 (P.D.A.), R01 AG081347 (W.D.), R01 AG069206 and (M.P.), R01 HL149992 (R.B.), R21 AG075446-02 (C.G.) and P30 AG068635 and R01 AG083373 (C.K.). H.T. was supported by a fellowship from the Uehara Memorial Foundation. T.M.M. was supported by a Conrad Prebys Foundation Predoctoral Fellowship. Sequencing was performed at the SBP Genomics Core by Brian James, Rebecca Porritt and Kang Liu, supported by P30 CA030199. This publication includes data generated at the UC San Diego IGM Genomics Center utilizing an Illumina NovaSeq 6000 purchased with funding from S10 OD026929. The sequencing analysis of human donor samples was supported by the Razavi Newman Integrative Genomics and Bioinformatics Core Facility of the Salk Institute (RRID:SCR_014842 and SCR_014846) with funding from P30 CA014195, P30 AG068635, P01 AG073084-04, the Howard and Maryam Newman Family Foundation and the Helmsley Trust. Single-cell sequencing was performed with assistance from Justin Buchanan and Allen Wang at the Single Cell Genomics Center for Epigenomics, UC San Diego. Compound preparation for stress assays was supported by Hilarie Austin and Atoosa Emami at the Conrad Prebys Center for Chemical Genomics High Throughput Screening Core, SBP, supported by P30 CA030199. Histology services were provided by Guillermina Garcia of the Histology Core, SBP, supported by P30 CA030199. AAV vector construction was provided by Chun-Teng Huang and Chih-Cheng Yang of the Functional Genomics Core, SBP, supported by P30 CA030199 and S10 OD036254. Flow cytometry support was provided by Yoav Altman of the Flow Cytometry Core, SBP, supported by P30 CA030199. Animal husbandry support and AAV injection were provided by Buddy Charbono and Adriana Charbono of the Animal Facility Core, SBP, supported by P30 CA030199. Manuscript writing and grammar checks were supported by ChatGPT (OpenAI).

---

**Extended Data Fig. 1.**
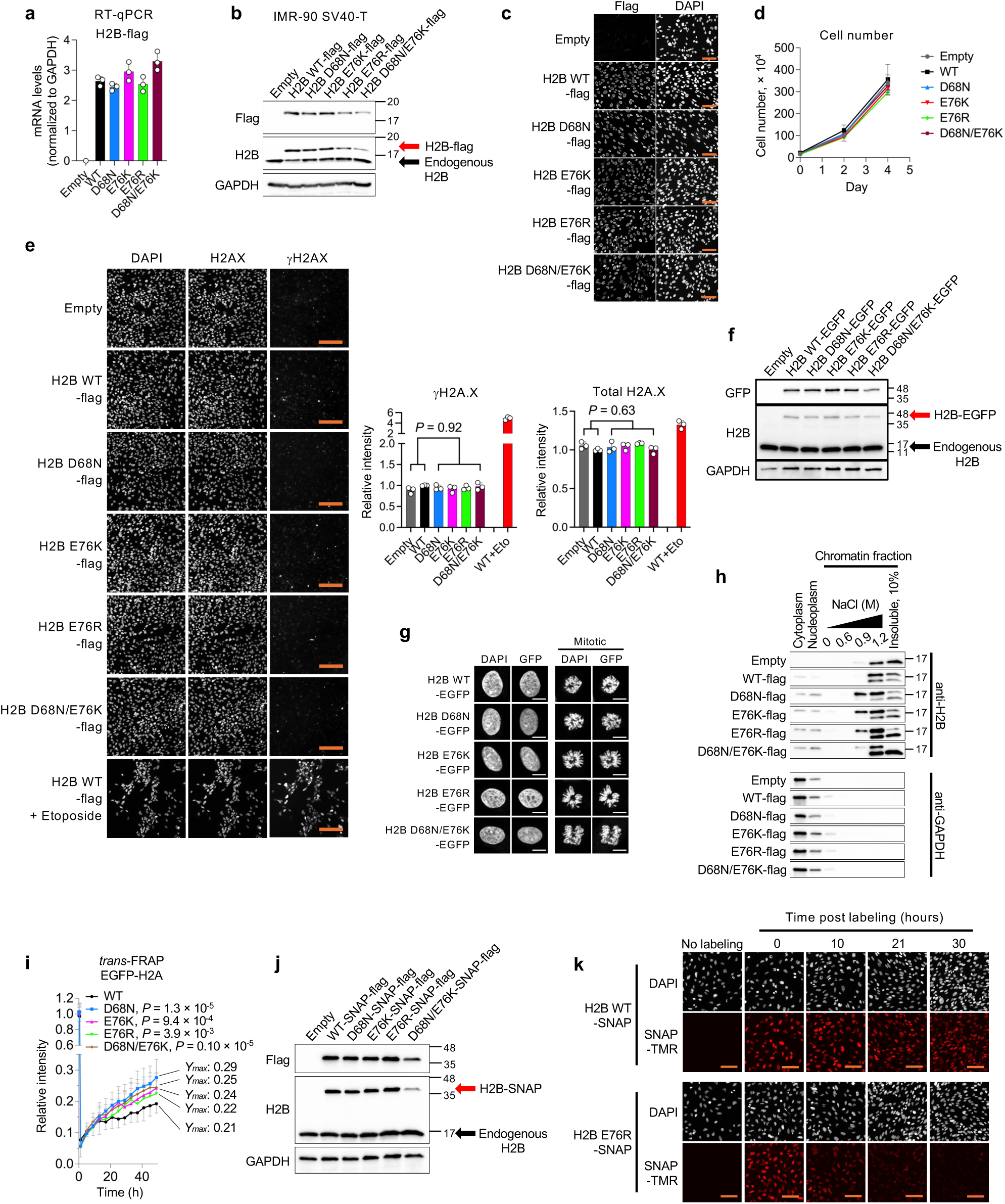
Effects of H2B mutants on nucleosome stability. **a**, RT-qPCR analysis of ectopically expressed WT or mutant H2B in IMR-90 SV40-T cells. Data represent mean ± s.d., normalized to GAPDH from three independent experiments. **b**, Western blot of ectopically expressed and endogenous H2B in IMR-90 SV40-T cells. GAPDH serves as a loading control. Data shown are representative of three independent experiments. **c**, Immunofluorescence images showing nuclear localization of Flag-tagged H2B WT or mutants. Scale bar, 100 μm. **d**, Cell proliferation over 4 days in IMR-90 SV40-T cells expressing WT or mutant H2B. Data represent mean ± s.d. from three independent wells per condition. **e**, Immunofluorescence images showing γH2AX and total H2AX in IMR-90 SV40-T cells expressing H2B WT, mutants or empty vector. Scale bar, 200 μm. Bar plots showing quantification of γH2AX and total H2AX intensity. As a positive control for double-stranded DNA breaks, cells were treated with etoposide at a final concentration of 10 μM for 24 h prior to fixation. Data represent mean ± s.d. from >1800 nuclei across three independent wells per condition. *P*-values were calculated using unpaired, two-tailed Student’s *t*-tests comparing Empty/WT and mutants. **f**, Western blot of EGFP-tagged H2B WT or mutants. GAPDH serves as a loading control. **g**, GFP fluorescence images of mitotic cells expressing GFP-tagged H2B WT or mutants. DNA was stained with DAPI. Scale bar, 10 μm. **h**, Sequential salt fractionation assay showing salt-dependent extraction of histones in cells expressing Flag-tagged H2B WT or mutants. Insoluble fraction was diluted 1:10 relative to other conditions. GAPDH was used as a cytoplasmic marker. **i**, Nucleosome stability assessed by FRAP in IMR-90 SV40-T cells expressing EGFP-tagged H2B WT or mutants. Data represent mean ± s.d. from >15 nuclei. The plateau value (*Ymax*) was calculated from the fitted recovery curve. *P*-values were calculated using unpaired, two-tailed Student’s *t*-tests comparing WT and each mutant at the final time point. **j**, Western blot of SNAP-tagged H2B WT or mutants. GAPDH serves as a loading control. **k**, Fluorescence images of SNAP-tagged H2B WT or mutants labeled with SNAP-TMR over a time course. Scale bar, 100 μm.

**Extended Data Fig. 2.**
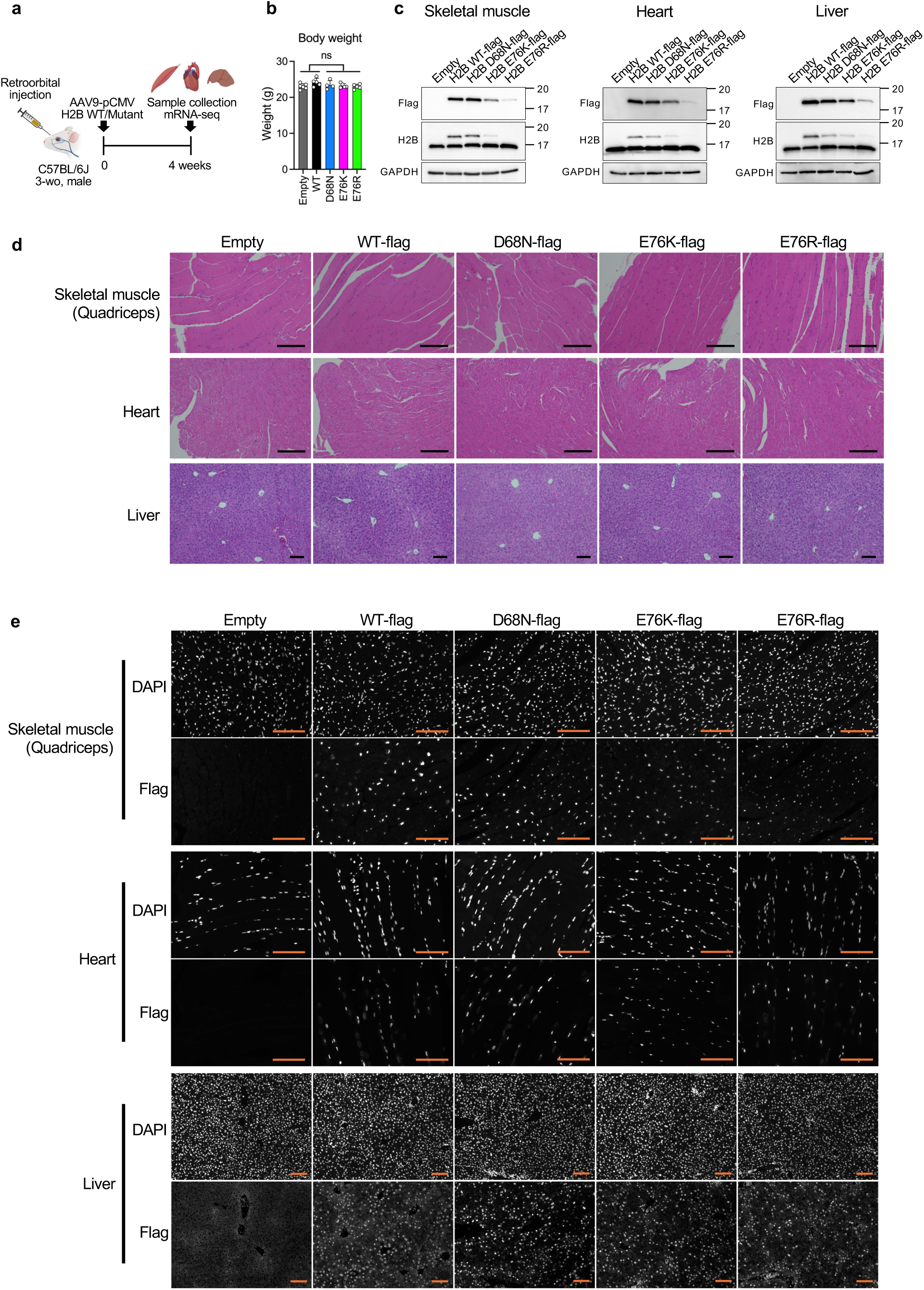
Transduction of mouse tissues using AAV9-pCMV mouse H2B WT, mutants, or an empty vector. **a**, Schematic of the experimental design. C57BL/6J male mice were retro-orbitally injected with AAV9-pCMV constructs expressing H2B WT, mutants (D68N, E76K, E76R) or an empty vector. Tissues were collected 4 weeks post-injection. **b**, Body weight measurements at the end of the experiment (n = 4–6 per group). *P*-values were calculated using unpaired, two-tailed Student’s *t*-tests comparing Empty/WT and mutants. **c**, Western blot of Flag-tagged H2B expression in mouse quadriceps skeletal muscle, heart and liver tissues. GAPDH serves as a loading control. **d**, H&E staining of skeletal muscle, heart and liver tissues from mice injected with H2B WT, mutant or empty constructs. Scale bars, 100 μm. **e**, Immunofluorescence images of Flag-tagged H2B WT or mutants in skeletal muscle, heart and liver tissues from mice injected with H2B WT, mutant or empty constructs. Scale bars, 100 μm.

**Extended Data Fig. 3.**
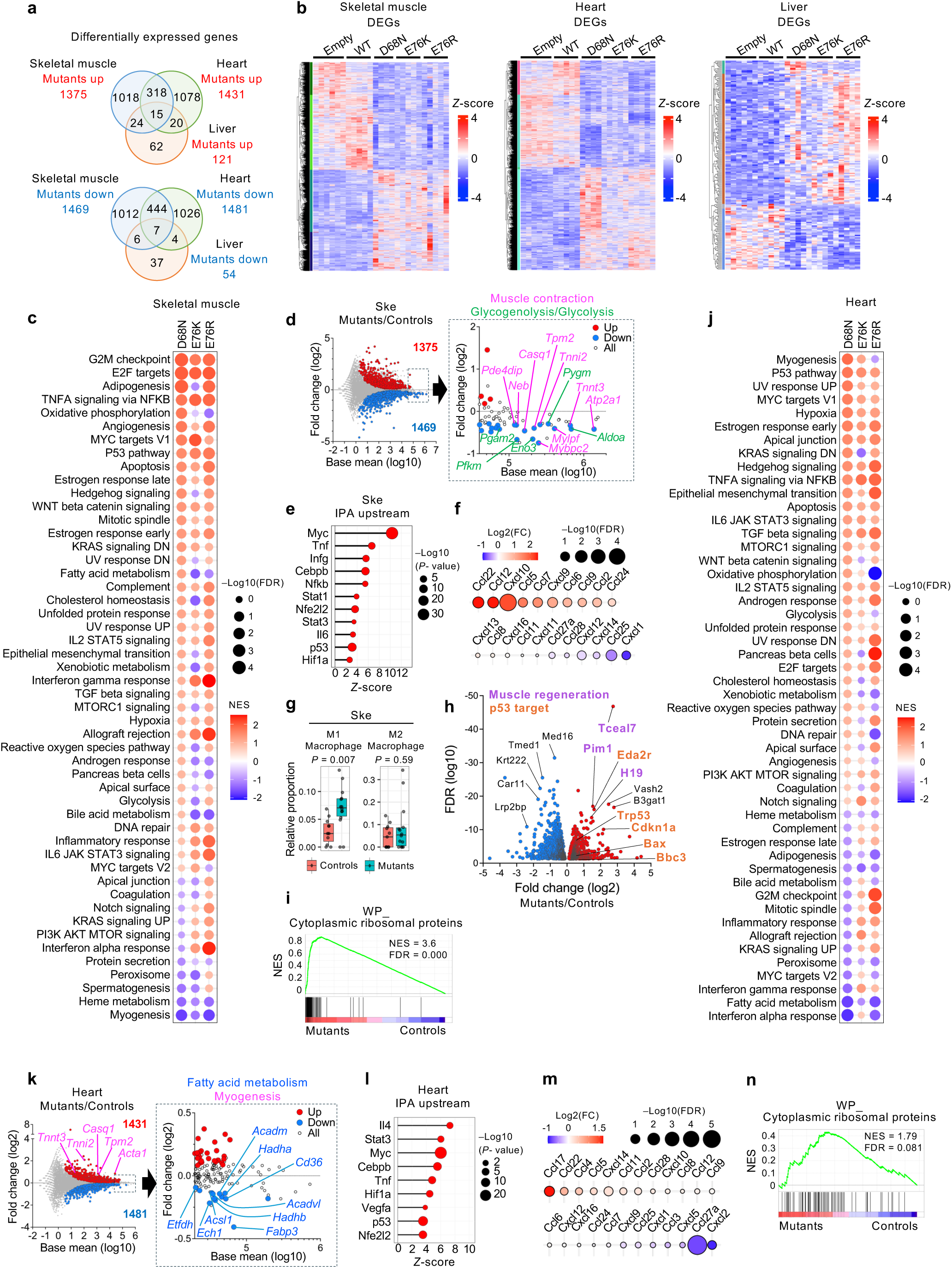
Effects of nucleosome instability on mouse tissues. **a**, Venn diagrams showing overlap of differentially expressed genes (DEGs) in skeletal muscle, heart and liver tissues from mice expressing H2B mutants compared to controls. **b**, Heatmaps of DEGs in skeletal muscle, heart and liver tissues. Red indicates upregulation; blue indicates downregulation. **c**, GSEA of hallmark pathways in transduced skeletal muscle. Dot color and size represent normalized enrichment score (NES) and FDR, respectively. **d**, MA plot of skeletal muscle comparing mutants to controls. Genes involved in myogenesis-related pathway such as muscle contraction and glycogenolysis/glycolysis are highlighted. **e**, IPA upstream regulator analysis showing activation of inflammatory and stress-response genes in mutant-expressing skeletal muscle. **f**, Relative expression fold-changes of chemokines in mutant-expressing skeletal muscle compared to Empty and WT. Dot color and size represent fold change and FDR, respectively. **g**, Box plot comparing proportions of M1 and M2 macrophage signatures in bulk mRNA-seq data between mutants and controls in skeletal muscle. **h**, Volcano plot of DEGs in mutant-expressing skeletal muscle compared to controls. Highlighted genes are associated with pathways related to muscle regeneration or p53 signaling. **i**, GSEA showing increased expression of cytoplasmic ribosomal protein genes in transduced skeletal muscle. **j**, GSEA of hallmark pathways in transduced heart. Dot color and size represent NES and FDR, respectively. **k**, MA plot of heart comparing mutants to controls. Genes involved in fatty acid metabolism and myogenesis are highlighted. **l**, IPA upstream regulator analysis showing activation of inflammatory and stress-response genes in mutant-expressing heart. **m**, Relative expression fold-changes of chemokines in mutant-expressing heart compared to Empty and WT. Dot color and size represent fold change and FDR, respectively. **n**, GSEA showing increased expression of cytoplasmic ribosomal protein genes in transduced heart.

**Extended Data Fig. 4.**
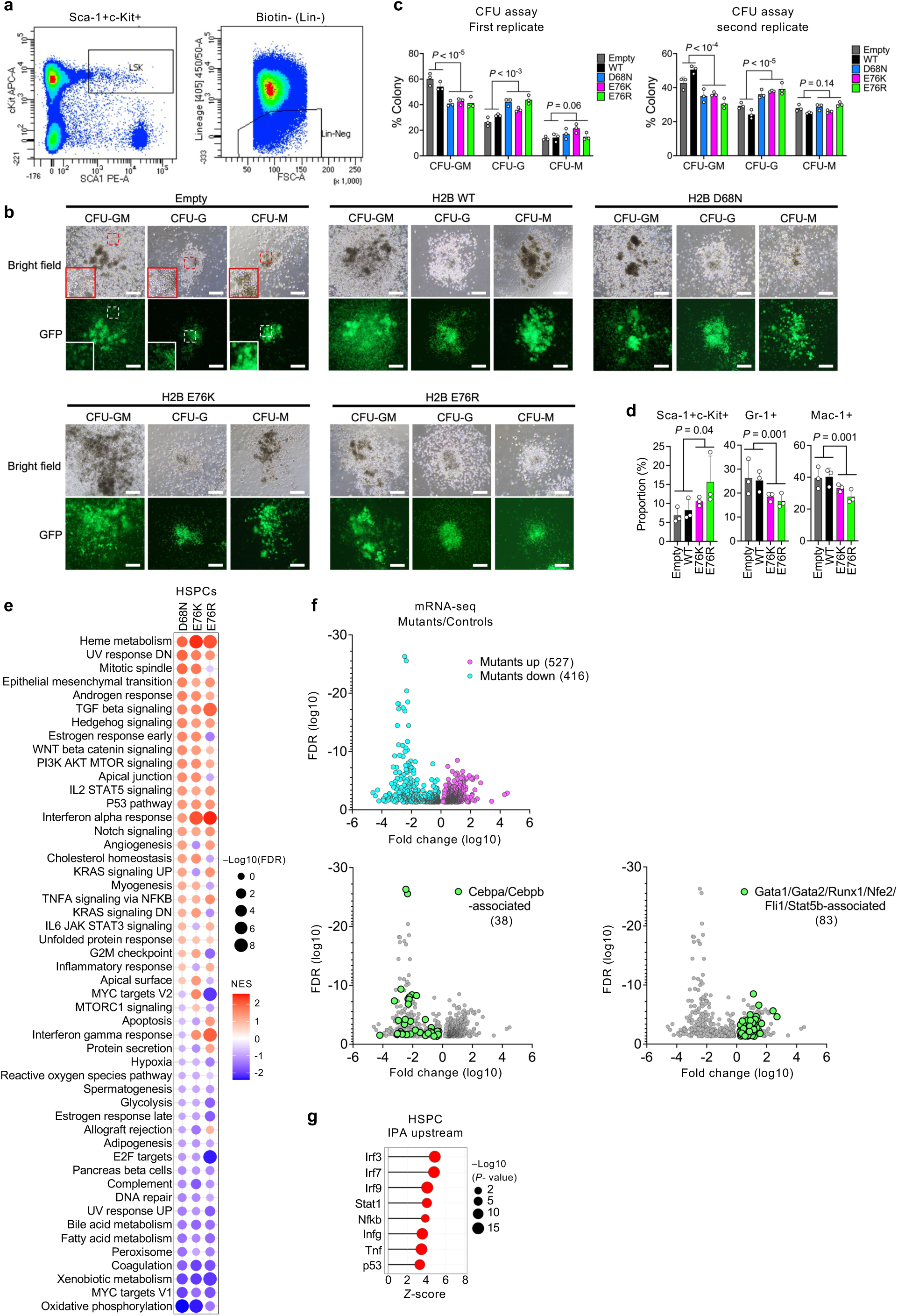
Effects of nucleosome instability on HSPC differentiation. **a**, Flow cytometry analysis of Lin⁻Sca-1⁺c-Kit⁺ populations from mouse bone marrow. **b**, Representative bright field and GFP fluorescence images from colony-forming unit (CFU) assays showing CFU-GM, CFU-G and CFU-M colonies derived from HSPCs transduced with GFP-marked empty vector, H2B WT or mutants after 8 days of differentiation. Scale bar, 200 μm. **c**, Quantification of CFU colony types from two independent replicates. Data represent mean ± s.d. for each condition. *P*-values were calculated using unpaired, two-tailed Student’s *t*-tests comparing Empty/WT and mutants. **d**, Recovery percentages of Sca-1⁺c-Kit⁺, Gr-1⁺, and Mac-1⁺ cells in cultures after 9 days of differentiation. *P*-values were calculated using unpaired, two-tailed Student’s *t*-tests comparing Empty/WT and mutants. **e**, GSEA of hallmark pathways in transduced skeletal muscle. Dot color and size represent normalized enrichment score (NES) and FDR, respectively. **f**, Volcano plots of DEGs in transduced HSPCs after 9 days of differentiation. Highlighted genes are associated with pathways relevant to myeloid and megakaryocyte/erythroid differentiation. **g**, IPA upstream regulator analysis showing activation of inflammatory and stress-response genes in mutant-expressing heart.

**Extended Data Fig. 5.**
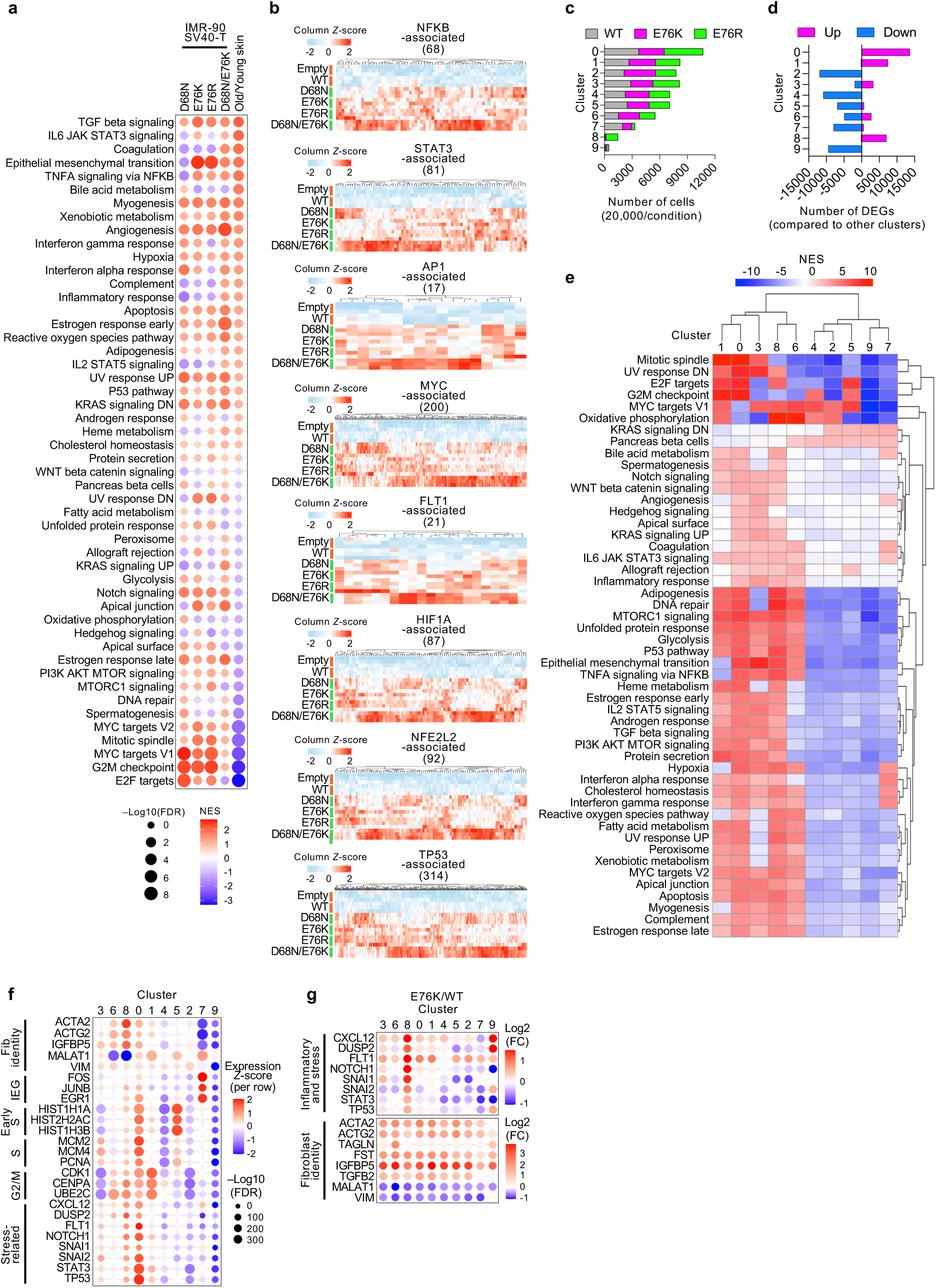
Effects of nucleosome instability on human fibroblasts. **a**, GSEA of hallmark pathways in mutant-expressing IMR-90 SV40-T cells compared to Empty and WT cells and in aged primary skin fibroblasts compared to young fibroblasts. **b**, Expression *Z*-scores of inflammatory and stress-response genes upregulated in mutant-expressing IMR-90 SV40-T cells compared to Empty and WT cells. **c**, Proportion of cells assigned to each of nine transcriptionally defined clusters in scRNA-seq of IMR-90 SV40-T WT, E76K or E76R cells. Cell numbers were standardized to 20,000 per condition. **d**, Number of significantly differentially expressed genes (FDR < 0.05) in each cluster compared to all other clusters in scRNA-seq analysis. **e**, GSEA of hallmark pathways in each cluster compared to all other clusters in scRNA-seq analysis. **f**, Relative expression *Z*-scores of fibroblast identity markers, cell-cycle genes and stress-related genes across clusters in scRNA-seq analysis. **g**, Cluster-wise log_2_ fold-changes in expression of inflammatory/stress-related or fibroblast identity genes in scRNA-seq analysis of E76K-versus WT-expressing IMR-90 SV40-T cells.

**Extended Data Fig. 6.**
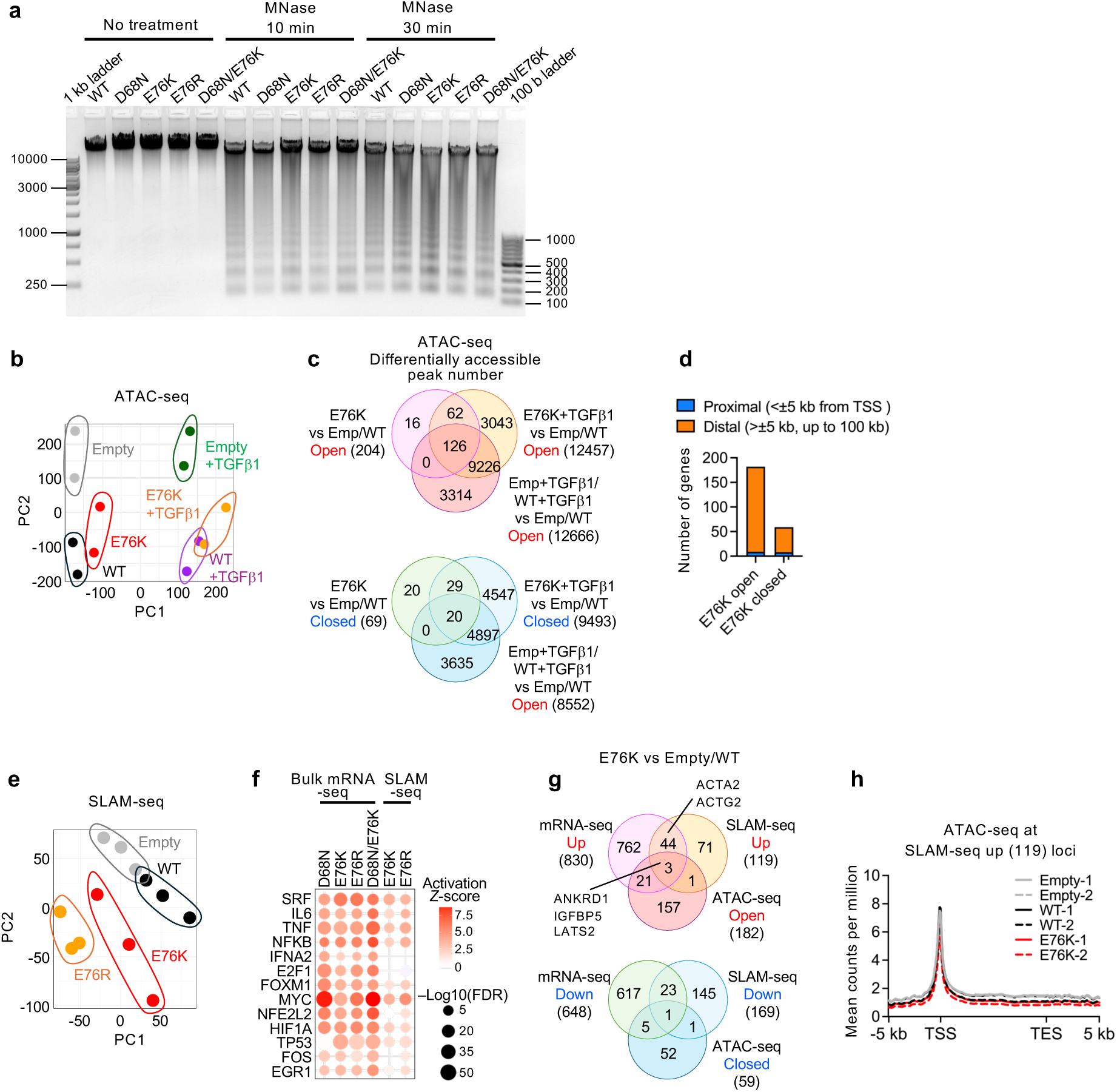
Effects of nucleosome instability on chromatin accessibility and nascent transcription. **a**, Gel electrophoresis showing chromatin digestion by MNase in IMR-90 SV40-T cells expressing H2B WT, mutants or empty vector. **b**, PCA of ATAC-seq data from IMR-90 SV40-T cells expressing H2B WT, E76K or empty vector, with or without TGFβ1 treatment (n = 2 per condition). **c**, Differentially accessible ATAC-seq peaks identified in IMR-90 SV40-T cells expressing H2B WT, E76K or empty vector, with or without TGFβ1 treatment. **d**, Number of genes associated with ATAC-seq peaks located at proximal (<5 kb from TSS) or distal (>25 kb, up to 100 kb) regions. Gene association was determined by proximal to TSS, extending up to 100 kb if no gene was found within 5 kb. **e**, PCA of SLAM-seq data from IMR-90 SV40-T cells expressing H2B WT, E76K, E76R or empty vector (n = 3 per condition). **f**, IPA upstream regulator analysis comparing bulk mRNA-seq and SLAM-seq data, indicating consistent activation of SRF-associated, inflammation and stress-response pathways in mutant-expressing cells compared to Empty and WT cells. **g**, Comparison across mRNA-seq, ATAC-seq and SLAM-seq analyses showing consistent upregulated/opened or downregulated/closed genes in E76K-expressing cells compared to Empty and WT cells. **h**, Meta-gene profile of ATAC-seq signals at gene loci upregulated in E76K-expressing cells on SLAM-seq analysis compared to Empty and WT cells.

**Extended Data Fig. 7.**
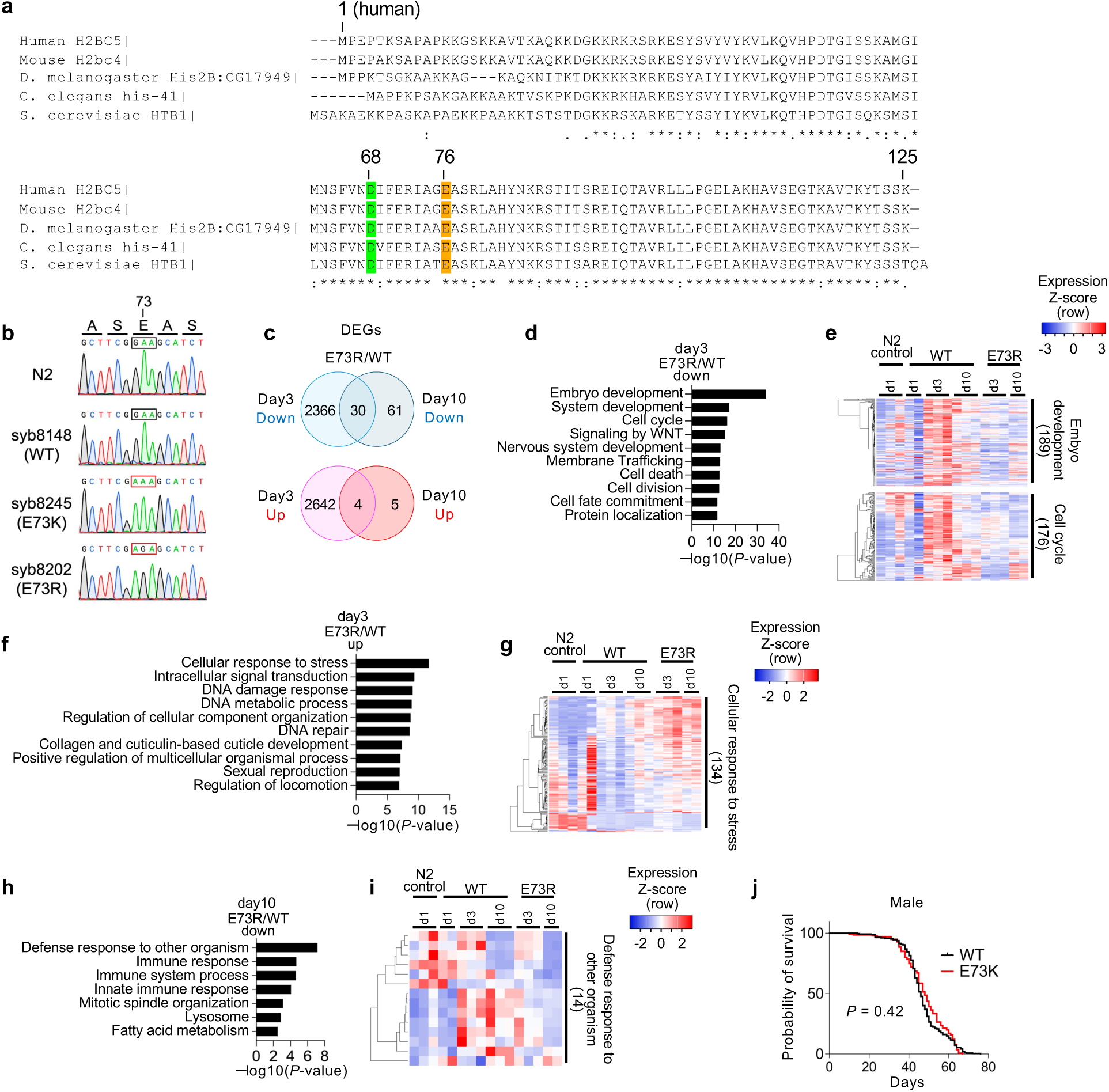
Effects of nucleosome instability on organismal aging. **a**, Sequence alignment of representative H2B genes from human, mouse, *D. melanogaster*, *C. elegans* and *S. cerevisiae*, highlighting conserved residues at D68 and E76 in human H2B. **b**, Sanger sequencing chromatograms confirming the wild-type sequence in N2 and WT strains and E73K and E73R mutations in *his-41* gene. **c**, Venn diagrams showing DEGs in bulk mRNA-seq of E73R mutants compared to WT at days 3 and 10 of adulthood. **d**, Gene ontology enrichment (Metascape) of 2,396 DEGs downregulated in E73R mutants at day3 post-adulthood compared to WT at day 3. **e**, Heatmaps showing expression of genes associated with embryo development and cell cycle, downregulated in E73R mutants at day 3 compared to WT at day 3. **f**, Gene ontology enrichment (Metascape) of 2,646 DEGs upregulated in E73R mutants at day 3 post-adulthood compared to WT at day 3. **g**, Heatmap of genes involved in cellular response to stress, upregulated in E73R mutants at day 3 compared to WT at day 3. **h**, Gene ontology enrichment (Metascape) of 91 DEGs downregulated in E73R mutants at day 10 post-adulthood compared to WT at day 10. **i**, Heatmap of genes related to defense response, upregulated in E73R mutants at day 10 compared to WT at day 10. **j**, Kaplan–Meier survival curves of male *D. melanogaster* expressing H2B WT or E73K. A total of 260 WT and 140 E73K flies were analyzed. *P*-value was calculated using the log-rank test comparing E73K to WT.

**Extended Data Fig. 8.**
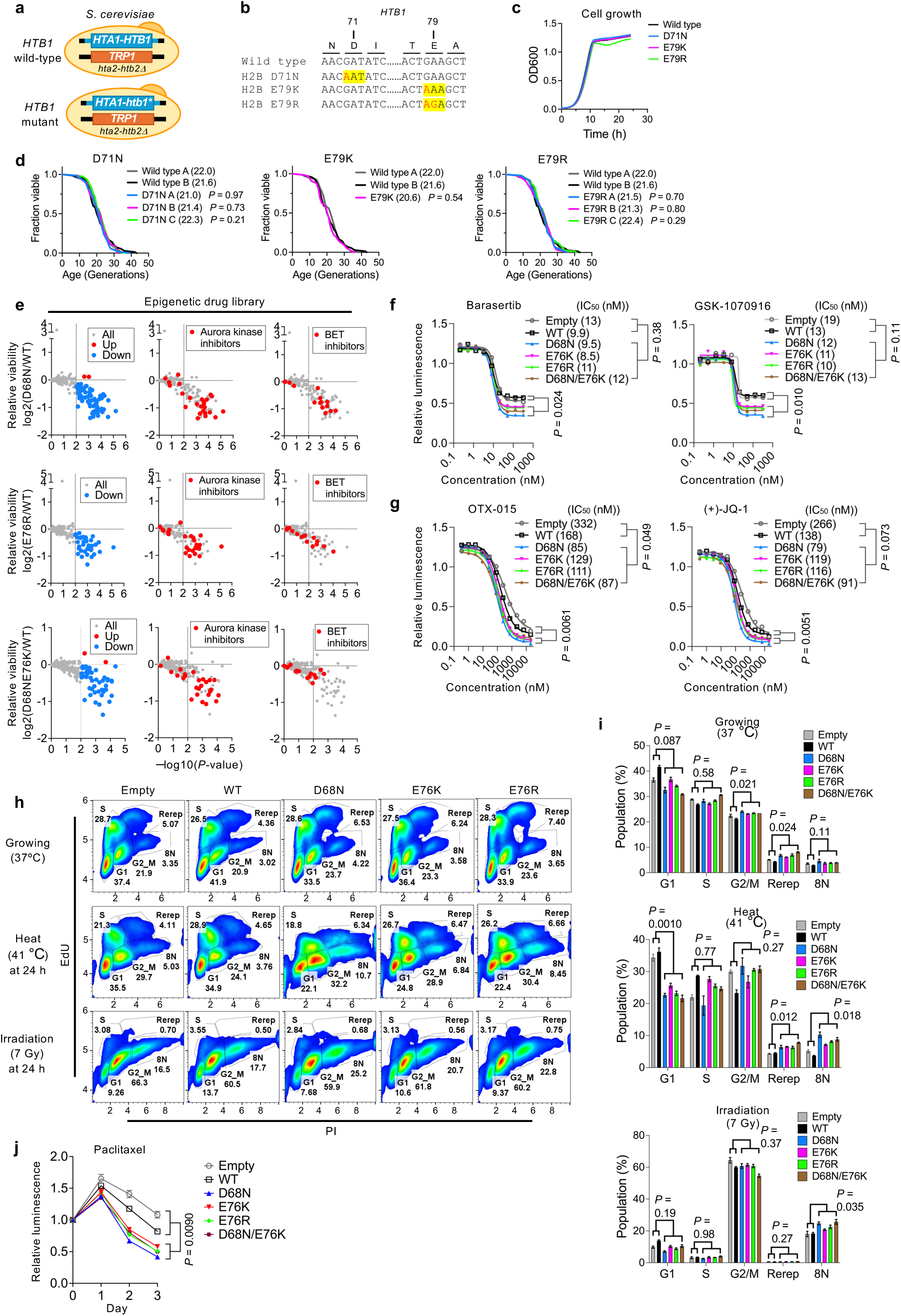
Effects of nucleosome instability on cellular stress resilience. **a**, Schematic of *S. cerevisiae* histone shuffle strains (FY406 background) expressing wild-type or mutant H2B/*HTB1*. In this strain, both endogenous H2A–H2B gene pairs are deleted, and viability is maintained by a plasmid carrying HTA1–HTB1, enabling targeted mutagenesis of HTB1. **b**, Sequence alignment of *HTB1* D71N, E79K and E79R mutants. **c**, Growth curves of wild-type and mutant yeast strains measured by OD600 over time. **d**, Kaplan–Meier survival curves of yeast expressing wild-type or mutant H2B. Log-rank tests were used to compare each mutant to WT; *P*-values are indicated on the graph. **e**, Relative cell viability of mutant-expressing IMR-90 SV40-T cells compared to WT-expressing cells across a library of 336 epigenetic compounds. Cell viability was measured on day 4 post-treatment. Data represent the relative fold-change of the means between each mutant and WT from three independent plates per condition. **f**,**g**, Dose–response curves for Barasertib and GSK-1070916 (Aurora kinase inhibitors) and OTX-015 and (+)-JQ-1 (BET inhibitors) in IMR-90 SV40-T cells expressing H2B WT, mutants or empty vector, with corresponding IC_50_ values. Data represent means from three replicates, each consisting of six wells per condition. *P*-values were calculated using unpaired, two-tailed Student’s *t*-tests on IC_50_ values and on the means of all mutants compared to Empty and WT at the bottom plateau from nonlinear regression. **h**, Flow cytometry analysis of cell cycle distribution 24 h after normal growth (37 °C), heat stress (41 °C) or irradiation (7 Gy). Data are representative of three independent replicates. Cell cycle phases (G1, S, G2/M) were determined by EdU incorporation and PI intensity. Populations with DNA content greater than 4N were also quantified to assess potential re-replication or polyploidy. **i**, Quantification of cell-cycle populations from h. *P*-values were calculated using unpaired, two-tailed Student’s *t*-tests comparing all mutants to Empty and WT. **j**, Cell viability assay with 0.01 μM paclitaxel in IMR-90 SV40-T cells expressing H2B WT, mutants or empty vector. Data represent mean ± s.d. from three replicates, each consisting of 16 wells per condition. *P*-values were calculated using unpaired, two-tailed Student’s *t*-tests comparing the means of all mutants to Empty and WT at the final time point.

**Supplementary Table. 1**: Demographics and experimental passage number of primary human dermal fibroblasts in RNA-seq experiment.

